# Reliance of neuronal gene expression on cohesin scales with chromatin loop length

**DOI:** 10.1101/2021.02.24.432639

**Authors:** Lesly Calderon, Felix D Weiss, Jonathan A Beagan, Marta S Oliveira, Yi-Fang Wang, Thomas Carroll, Gopuraja Dharmalingam, Wanfeng Gong, Kyoko Tossell, Vincenzo de Paola, Chad Whilding, Mark A. Ungless, Amanda G Fisher, Jennifer E Phillips-Cremins, Matthias Merkenschlager

**Affiliations:** MRC London Institute of Medical Sciences, Institute of Clinical Sciences, Faculty of Medicine, Imperial College London, Du Cane Road, London W12 0NN, UK; Institute of Clinical Sciences, Faculty of Medicine, Imperial College London, Du Cane Road, London W12 0NN, UK; Department of Bioengineering, University of Pennsylvania, Philadelphia, PA 19104, USA; Epigenetics Program, Perelman School of Medicine, University of Pennsylvania, Philadelphia, PA 19104, USA; Department of Genetics, Perelman School of Medicine, University of Pennsylvania, Philadelphia, PA 19104, USA

**Author notes:** Equal contribution (Lesly Calderon, Felix D Weiss, Jonathan A Beagan). IMP, Vienna, Austria (LC), Institute of Innate Immunity, University of Bonn, Germany (FDW) Yale University, New Haven, CT, USA (JAB), Rockefeller University, NY, USA (TC), Department of Life Sciences, Imperial College London, UK (KT) Mental Health Innovations, London, UK (MAU).

## Abstract

Cohesin and CTCF are major drivers of 3D genome organization, but their role in neurons is still emerging. Here we show a prominent role for cohesin in the expression of genes that facilitate neuronal maturation and homeostasis. Unexpectedly, we observed two major classes of activity-regulated genes with distinct reliance on cohesin in primary cortical neurons. Immediate early genes remained fully inducible by KCl and BDNF, and short-range enhancer-promoter contacts at the Immediate early gene *Fos* formed robustly in the absence of cohesin. In contrast, cohesin was required for full expression of a subset of secondary response genes characterised by long-range chromatin contacts. Cohesin-dependence of constitutive neuronal genes with key functions in synaptic transmission and neurotransmitter signaling also scaled with chromatin loop length. Our data demonstrate that key genes required for the maturation and activation of primary cortical neurons depend cohesin for their full expression, and that the degree to which these genes rely on cohesin scales with the genomic distance traversed by their chromatin contacts.

## Introduction

Mutations in cohesin and CTCF cause intellectual disability in humans (Deardorff et al., 2018; Gregor et al., 2013; Rajarajan et al., 2016), and defects in neuronal gene expression (Kawauchi *et al*. 2009; Fujita et al., 2017; van den Berg *et al*., 2017; McGill et al., 2018; Yamada et al., 2019; Weiss et al., 2021), neuronal morphology (Fujita et al., 2017; McGill et al., 2018; Sams et al., 2016), long-term potentiation (Sams et al., 2016; Kim et al., 2018), learning, and memory (McGill et al., 2018; Yamada et al., 2019; Sams et al., 2016; Kim et al., 2018) in animal models. In addition to mediating canonical functions in the cell cycle (Nasmyth & Haering, 2009), cohesin cooperates with CTCF to facilitate the spatial organisation of the genome in the nucleus. Cohesin traverses chromosomal DNA in an ATP-dependent manner through a mechanism known as loop extrusion. This process generates self-interacting domains that are delimited by chromatin boundaries marked by CTCF and defined by an increased probability of chromatin contacts (Fudenberg et al., 2016; Rao et al., 2014; Rao et al., 2017; Nora et al. 2017; Schwarzer et al., 2017). The resulting organisation of the genome into domains and loops is thought to contribute to the regulation of gene expression by facilitating appropriate enhancer-promoter interactions (Dekker & Mirny, 2016; Merkenschlager & Nora, 2016; Beagan & Phillips-Cremins, 2020; McCord et al., 2020) and its disruption can cause human disease (Lupiáñez et al. 2015; Spielmann et al., 2018; Sun et al., 2018), including neurodevelopmental disorders (Won et al., 2016).

Recent studies which induced global cohesin loss on acute timescales resulted in the altered expression of only a small number of genes in a human cell line in vitro (Rao et al., 2017), while later time points after cohesin depletion in vivo revealed more pervasive disruption in expression (Schwarzer et al., 2017). We recently reported that cohesin loss has modest effects on constitutive gene expression in uninduced macrophages, but severely disrupted the establishment of new gene expression programmes upon the induction of a new macrophage state (Cuartero et al., 2018). These data support a model in which cohesin-mediated loop extrusion is more important for the establishment of new gene expression rather than maintenance of existing programs (Cuartero et al., 2018). However, the applicability of this model across other cell types remains unclear, and the extent to which deficits in cohesin function alter neuronal gene expression remains a critical underexplored question.

Activity-regulated neuronal genes (ARGs) are defined by transcriptional induction in response to neuronal activity, and are important for cellular morphology, the formation of synapses and circuits, and ultimately for learning and memory (Gallo et al., 2018; Kim et al., 2010; Malik et al., 2014; Greer et al., 2008; Tyssowski et al., 2018; Yap et al., 2018). ARG induction is accompanied by acetylation of H3K27, as well as the recruitment of RNA polymerase 2, cohesin, and other chromatin binding proteins at ARGs and their enhancers (Greer et al., 2008; Malik et al., 2014; Yamada et al., 2019; Yap et al., 2018; Tyssowski et al., 2018; Schaukowitch et al., 2014; Beagan et al., 2020). For a subset of ARGs, long-range contacts between enhancers and promoters increase upon stimulation of post-mitotic neurons (Schaukowitch et al., 2014; Sams et al., 2016; Beagan et al., 2020). Among neuronal ARGs, immediate early genes (IEGs) and secondary response genes (SRGs) are known to be activated on different time scales according to distinct mechanisms (Tyssowski et al., 2018). Moreover, IEGs and SRGs differ significantly in their looping landscape, as IEGs form fewer, shorter enhancer-promoter contacts compared to the complex, long-range interactions formed by many SRGs (Beagan et al., 2020). Given the importance of cohesin and regulatory looping interactions for brain function, there is a strong imperative to understand the functional role of cohesin-dependent loops in the establishment and maintenance of developmentally-regulated and activity-stimulated neuronal gene expression programs.

Here we establish an experimental system to address the role of cohesin in 3D genome organisation and gene expression in non-dividing, post-mitotic neurons, independently of essential cohesin functions in the cell cycle. We employ developmentally regulated *Nex*^Cre^ expression (Goebbels et al., 2006; Hirayama et al., 2012) to inducibly deplete the cohesin subunit RAD21 specifically in immature post-mitotic mouse neurons *in vivo*. Cohesin depletion in immature post-mitotic mouse neurons disrupted CTCF-based chromatin loops, and reduced the expression of neuronal genes related to synaptic transmission, neuronal development, adhesion, connectivity, and signaling, resulting in impaired neuronal maturation. Neuronal ARGs were pervasively deregulated in cohesin-deficient neurons, consistent with a model in which the establishment of new gene expression programs in both macrophages and neurons is driven in part by cohesin-mediated enhancer-promoter contacts (Rajarajan et al., 2016; Yamada et al., 2019; Sams et al., 2016; Schaukowitch et al., 2014; Cuartero et al., 2018). The cohesin-dependence of ARGs was confirmed in an inducible system that allows for proteolytic cohesin cleavage in primary neurons (Weiss et al., 2021), providing temporal control over cohesin levels on a time scale similar to the establishment of neuronal gene expression programs upon neural stimulation, thus allowing us to disentangle the role for cohesin-mediated chromatin contacts in the maintenance of existing transcriptional programmes versus the establishment of new transcriptional programs in neural circuits. Surprisingly, despite pervasive deregulation at beaseline, most IEGs and a subset of SRGs remained fully inducible by KCl and BDNF stimulation after genetic or proteolytic depletion of cohesin.

Cohesin-dependent and -independent ARGs were distinguished not by the binding of cohesin or CTCF to their promoters (Schaukowitch et al., 2014; Sams et al., 2016), but instead by their 3D connectivity. SRGs that depended on cohesin for full inducibility engaged in longer chromatin loops than IEGs, or SRGs that remained fully inducible in the absence of cohesin. Unlike ARGs, the majority of neuronal genes that mediate synaptic transmission and neurotransmitter signaling are constitutively expressed. Nevertheless, as with ARGs, the reliance of these key neuronal genes on cohesin scaled with chromatin loop length.

Consistent with a model where short-range enhancer-promoter loops can form in the absence of cohesin, we find that the enhancer activity and short-range enhancer-promoter contacts at the IEG *Fos* remained robustly inducible in cohesin-depleted neurons. Finally, re-expression of RAD21 protein in cohesin-depleted neurons re-established lost chromatin loops and restored wild-type expression levels of disrupted SRGs, and of constitutive neuronal genes engaged in long-range loops. Together, our data support a model where key neuronal genes required for the maturation and activation of primary neurons require cohesin for their full expression. The degree to which neuronal genes rely on cohesin scales with the genomic distance traversed by their chromatin loops, including loops connecting promoters with activity-induced enhancers.

## Results

### Conditional deletion of cohesin in immature post-mitotic neurons

To explore the role of cohesin in post-mitotic neurons, we deleted the essential cohesin subunit RAD21 (*Rad21*^lox^, Seitan et al., 2011) in immature cortical and hippocampal neurons using *Nex*^Cre^ (Goebbels et al., 2006; Hirayama et al., 2012). Explant cultures of wild-type and *Rad21*^lox/lox^ *Nex*^Cre^ E17.5/18.5 cortex contained >95% MAP2^+^ neurons, with <1% GFAP^+^ astrocytes or IBA1^+^ microglia (Fig. 1a). Immunofluorescence staining showed the loss of RAD21 protein specifically in GAD67^-^ neurons (Fig. 1b, c). This was expected, as *Nex*^Cre^ is expressed by excitatory but not by inhibitory neurons (Goebbels et al., 2006; Hirayama et al., 2012). Consistent with the presence of ∼80% of GAD67^-^ and ∼20% GAD67^+^ neurons in the explant cultures, *Rad21* mRNA expression was reduced by 75-80% overall (Fig. 1d, left). There was a corresponding reduction in RAD21 protein (Fig. 1d, middle). To focus our analysis on cohesin-deficient neurons we combined *Nex*^Cre^-mediated deletion of *Rad21* with *Nex*^Cre^-dependent expression of an epitope-tagged ribosomal subunit (*Rpl22*-HA RiboTag; Sanz et al., 2009). We verified that *Nex*^Cre^-induced RPL22-HA expression was restricted to RAD21-depleted neurons (Supplementary Fig. 1a) and performed high throughput sequencing of *Rpl22*-HA RiboTag-associated mRNA (RiboTag RNA-seq, Supplementary Data 1). Comparison with total RNA-seq (Supplementary Data 2) showed that *Nex*^Cre^ RiboTag RNA-seq captured excitatory neuron-specific transcripts, such as *Slc17a7* and *Camk2a.* Transcripts selectively expressed in astrocytes (*Gfap, Aqp4, Mlc1*), microglia (*Aif1*) and inhibitory neurons (*Gad1, Gad2, Slc32a1*) were depleted from *Nex*^Cre^ RiboTag RNA-seq (Supplementary Fig. 1b). RiboTag RNA-seq enabled an accurate estimate of residual *Rad21* mRNA, which was <5% in *Rad21*^lox/lox^ *Nex*^Cre^ cortical neurons (Fig. 1d, right). These data show near-complete loss of *Rad21* mRNA and undetectable levels of RAD21 protein in *Nex*^Cre^-expressing *Rad21*^lox/lox^ neurons. Analysis of chromatin conformation by Chromosome-Conformation-Capture-Carbon-Copy (5C) showed a substantial reduction in the strength of CTCF-based chromatin loops after genetic (*Rad21*^lox/lox^ *Nex*^Cre^, Fig. 1e) and after proteolytic cohesin depletion (RAD21-TEV, Supplementary Fig. 2a,b,c; Weiss et al., 2021). Of note, restoration of RAD21 expression rescued CTCF-based chromatin loop formation (Supplementary Fig. 2b,c). These data show that cohesin is directly linked to the strength of CTCF-based chromatin loops in primary in post-mitotic neurons.

**Figure 1.**
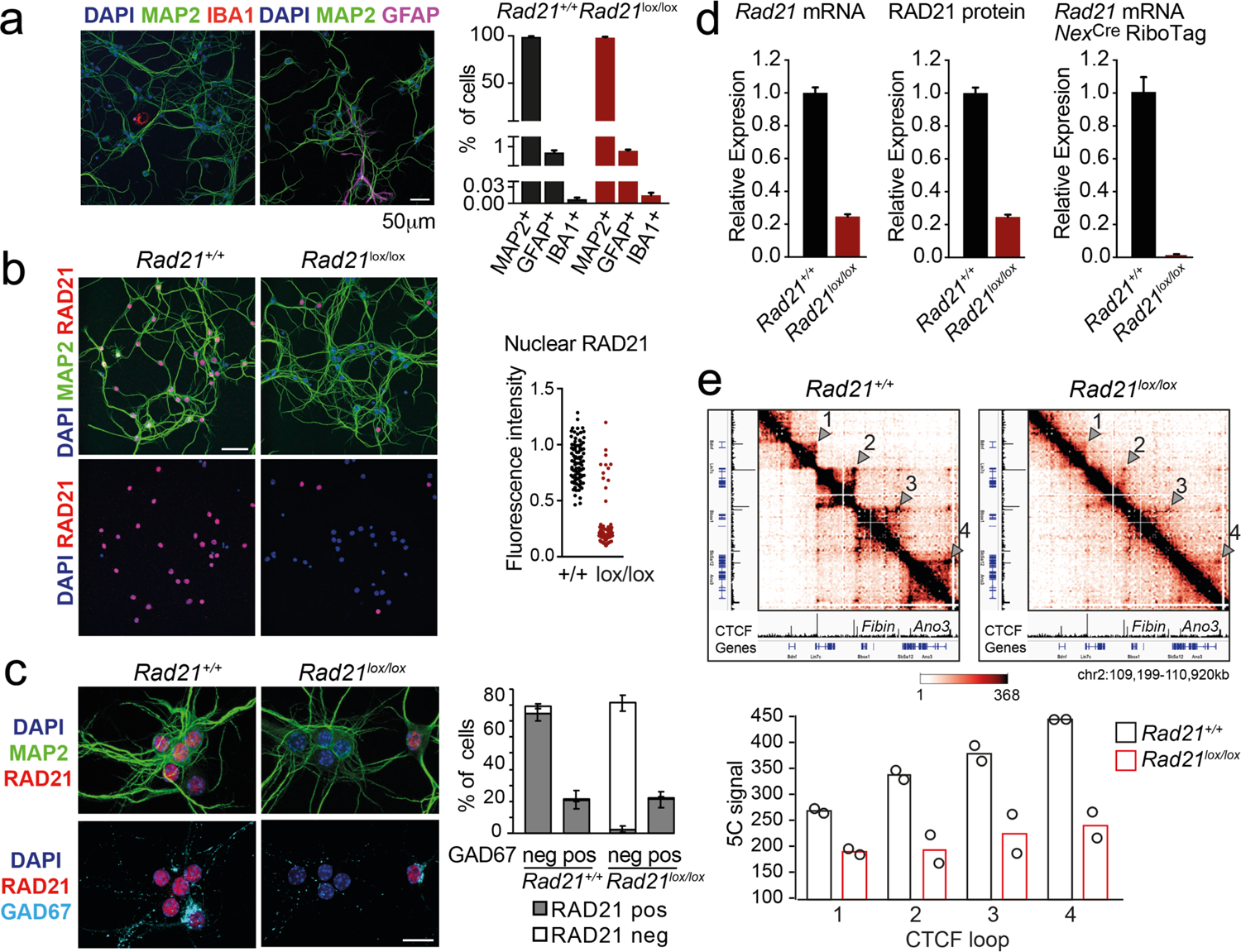
Conditional cohesin deletion in post-mitotic neurons. a) E17.5-E18.5 cortices were dissociated and plated on poly-D-lysine. After 10d, cultures were stained for pan neuronal (MAP2), astrocyte (GFAP) and microglia (IBA1) markers, and cell type composition was determined by quantitative analysis of immunofluorescence images. Based on 6 *Rad21^+/+^ Nex*^Cre^ and 8 *Rad21*^lox/lox^ *Nex*^Cre^ different samples analysed in 4 independent experiments. b) Immunofluorescence staining of *Rad21*^+/+^ *Nex*^Cre^ and *Rad21*^lox/lox^ *Nex*^Cre^ neuronal explant cultures for RAD21 and MAP2 (left) and distribution of RAD21 expression by MAP^+^ neurons (right). Note the discontinuous distribution of RAD21 expression in *Rad21*^lox/lox^ *Nex*^Cre^ neurons. Three independent experiments per genotype. DAPI marks nuclei. Scale bar = 60 μm. c) Immunofluorescence staining for RAD21, MAP2, and the marker of GABAergic inhibitory neurons, GAD67 (left). Distribution of RAD21 expression in GAD67^+^ and GAD67^-^ neurons (right). Note that the discontinuous distribution of RAD21 expression in *Rad21*^lox/lox^ *Nex*^Cre^ neuronal explant cultures is due to GAD67^+^ GABAergic inhibitory neurons. Three independent experiments for *Rad21^+/+^ Nex*^Cre^ and 6 independent experiments for *Rad21*^lox/lox^ *Nex*^Cre^. DAPI marks nuclei. Scale bar = 20 μm. d) Quantitative RT-PCR analysis of *Rad21* mRNA expression in *Rad21^+/+^ Nex*^Cre^ and *Rad21*^lox/lox^ *Nex*^Cre^ cortical explant cultures (mean ± SEM, n=18). *Hprt* and *Ubc* were used for normalization (left). RAD21 protein expression in *Rad21^+/+^ Nex*^Cre^ and *Rad21*^lox/lox^ *Nex*^Cre^ cortical explant cultures was quantified by fluorescent immunoblots (mean ± SEM, n=6, a representative blot is shown in Supplementary Fig. 1a) and normalised to LaminB (center). *Nex*^Cre^ RiboTag RNA-seq of analysis of *Rad21* mRNA expression in *Rad21^+/+^ Nex*^Cre^ and *Rad21*^lox/lox^ *Nex*^Cre^ cortical explant cultures (right, 3 independent biological replicates). e) 5C heat maps of a 1.72 Mb region on chromosome 2, comparing *Rad21^+/+^ Nex*^Cre^ and *Rad21*^lox/lox^ *Nex*^Cre^ cortical explant cultures. CTCF ChIP-seq in cortical neurons (Bonev et al., 2017) and mm9 coordinates are shown for reference. Arrowheads mark the position of CTCF-based loops. Results were consistent across two replicates and 3 chromosomal regions Histograms below show the quantification of representative CTCF-based loops (arrowheads) in two independent biological replicates for control and *Rad21*^lox/lox^ *Nex*^Cre^ neurons.

### Loss of cohesin from immature post-mitotic neurons perturbs neuronal gene expression

Using RiboTag RNA-seq to profile gene expression specifically in cohesin-depleted neurons we identified 1028 downregulated and 572 upregulated transcripts in *Rad21*^lox/lox^ *Nex*^Cre^ cortical neurons (Fig. 2a, Supplementary Fig. 3), with preferential deregulation of neuron-specific genes (*P* < 2.2e-16, see methods). Gene ontology (Fig. 2b) and gene set enrichment analysis (GSEA, Supplementary Fig. 3b) showed that downregulated genes in *Rad21*^lox/lox^ *Nex*^Cre^ neurons were enriched for synaptic transmission, neuronal development, adhesion, connectivity, and signaling (Supplementary Data 3) and showed significant overlap with genes linked to human ASD (*P* = 5.10E-15, Supplementary Fig. 3c; Banerjee-Basu & Packer, 2010). Upregulated genes showed no comparable functional enrichment (Fig. 2b). ARGs such as *Fos, Arc,* and *Bdnf* are activity-regulated by definition, and hence lowly expressed in basal conditions due to spontaneous synaptic activity. In *Rad21*^lox/lox^ *Nex*^Cre^ neurons, previously defined ARGs (Kim et al., 2010) were more frequently downregulated than constitutively expressed genes (Fig. 2c, *P* = 1.03e-16, odds ratio = 3.27). These data show that immature post-mitotic neurons require cohesin to establish and/or maintain the correct level of expression of genes that support neuronal maturation, including the growth and guidance of axons, the development of dendrites and spines, and the assembly, function, and plasticity of synapses.

**Figure 2.**
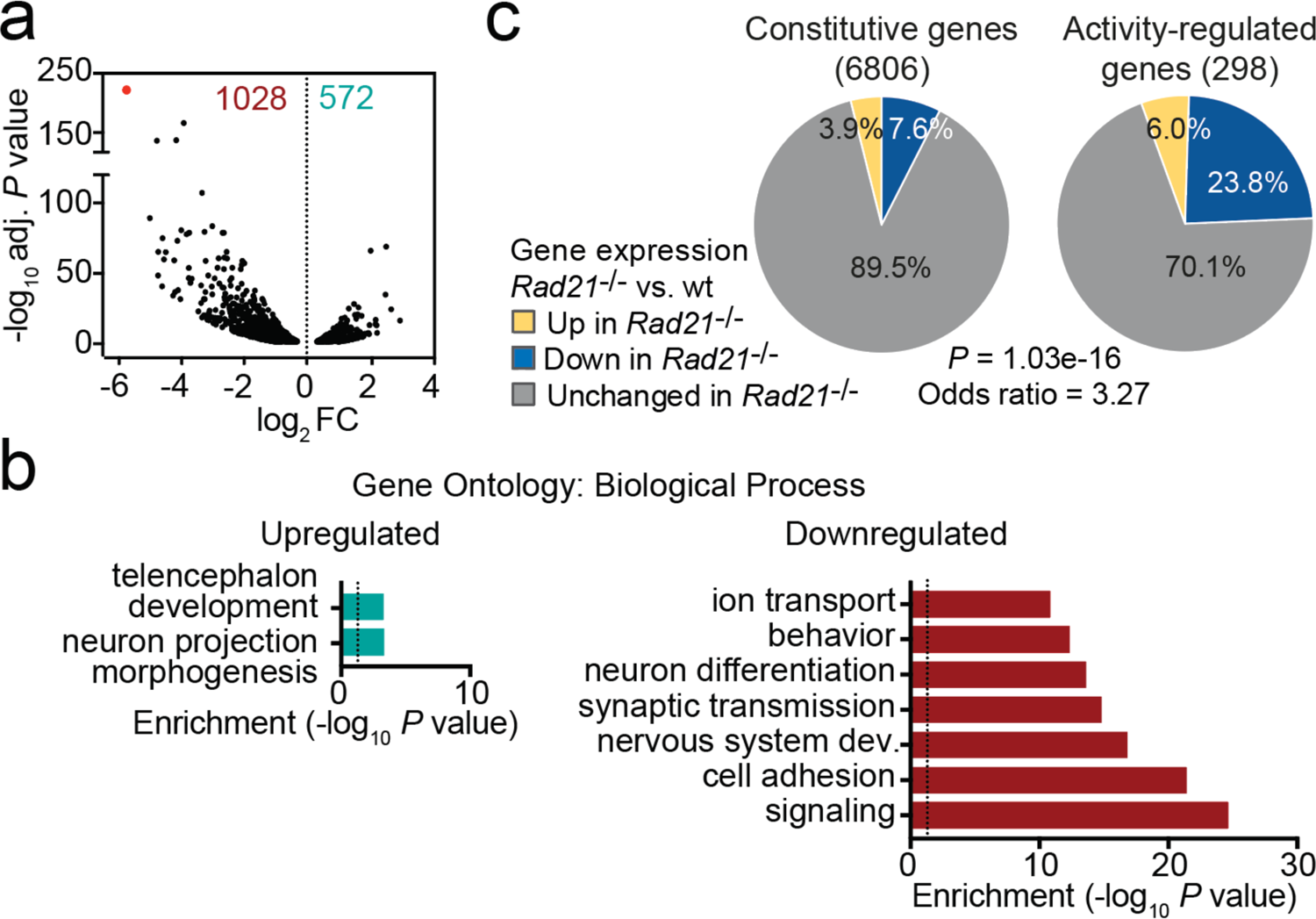
Loss of cohesin from immature post-mitotic neurons perturbs neuronal gene expression. a) Volcano plot representing log2 fold-change (FC) versus significance (-log10 of adjusted P values) of downregulated genes (1028) and upregulated genes (572) in RiboTag RNA-seq of *Rad21*^lox/lox^ *Nex*^Cre^ versus *Rad21^+/+^ Nex*^Cre^ neurons (Supplementary Data 1). Red marks *Rad21*. b) Analysis of gene ontology of biological functions of deregulated genes in *Rad21*^lox/lox^ *Nex*^Cre^ neurons. Enrichment is calculated relative to expressed genes (Supplementary Data 3). c) The percentage of constitutive (adj. *P* > 0.05 in KCl 1h vs TTX and KCl 6h vs TTX, see methods) and activity-regulated genes (Kim et al., 2010) found deregulated in *Rad21*^lox/lox^ *Nex*^Cre^ neurons in explant culture at baseline as determined by RiboTag RNA-seq. The *P*-value (Fisher Exact Test) and Odds ratio indicate that ARGs are more frequently deregulated than constitutive genes.

### A role for cohesin in the maturation of post-mitotic neurons

We next set out to examine the impact of cohesin deletion on the maturation of immature postmitotic neurons. *Rad21*^lox/lox^ *Nex*^Cre^ embryos were found at the expected Mendelian ratios throughout gestation, however postnatal lethality was evident (Supplementary Fig. 4a). *Rad21*^lox/lox^ *Nex*^Cre^ cortical neurons did not show increased proliferation (Supplementary Fig. 4b), no upregulation of apoptosis markers or signs of DNA damage (Supplementary Fig. 4b) and no stress-related gene expression (Supplementary Fig. 4c). Brain weight (Supplementary Fig. 4d) and cellularity (Supplementary Fig. 4e) were comparable between *Rad21*^+/+^ and *Rad21*^lox/lox^ *Nex*^Cre^ neocortex. The neuronal transcription factors TBR1, CTIP2 and CUX1 were expressed beyond the boundaries of their expected layers in *Rad21*^lox/lox^ *Nex*^Cre^ cortices, and deeper cortical layers appeared disorganised (Fig. 3a). These findings are consistent with the reported cohesin-dependence of neuronal guidance molecule expression (Kawauchi *et al*. 2009; Remeseiro *et al*. 2012; Guo *et al*. 2015) and migration (van den Berg *et al*., 2017). To assess the impact of cohesin on morphological maturation, we cultured cortical neurons in the presence of wild type glia (Kaech & Banker, 2006).

**Figure 3.**
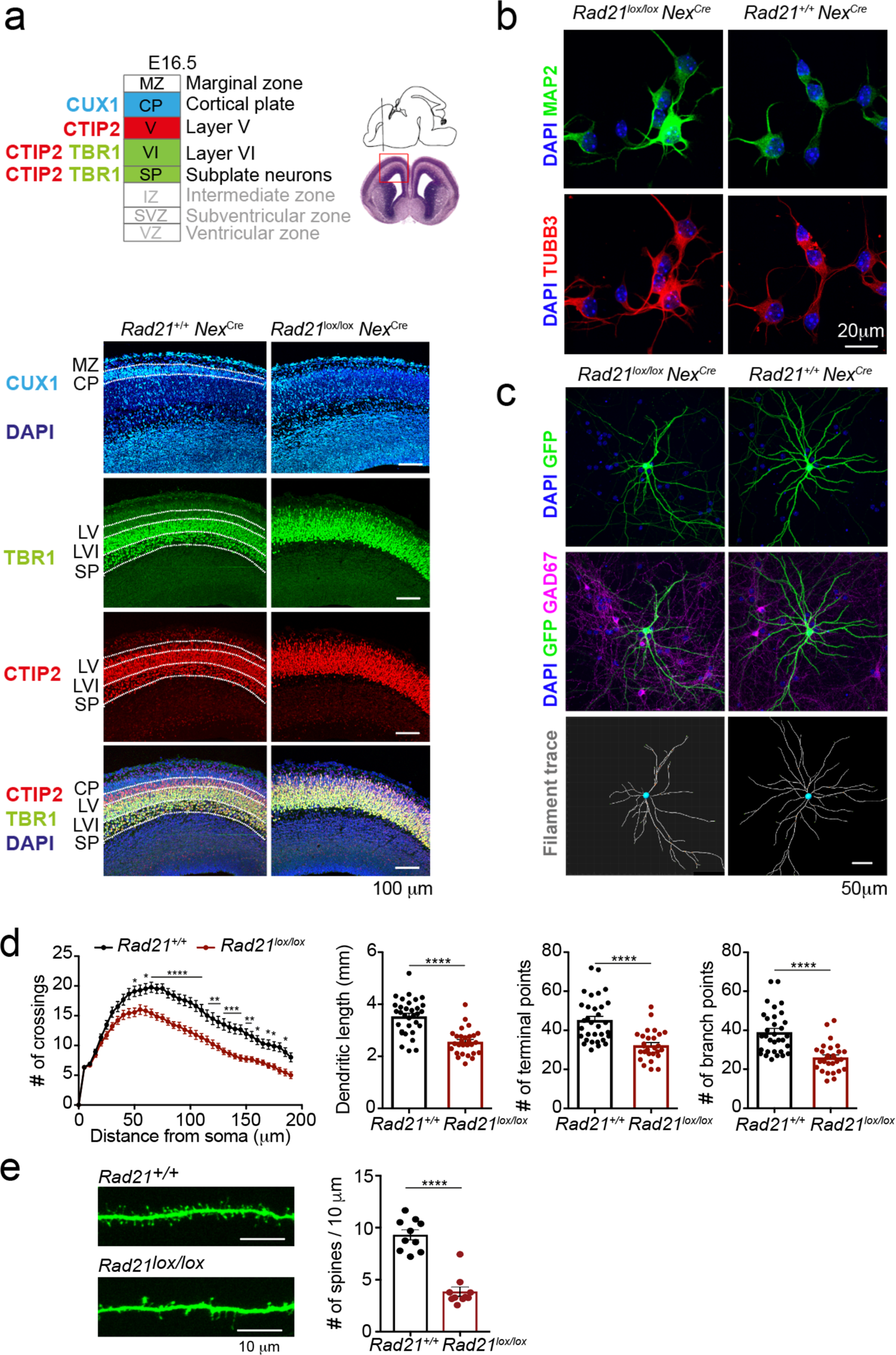
Cohesin contributes to the maturation of post-mitotic neurons. a) Schema of cortical layers (Greig et al., 2013) showing subplate (SP), layer 6 (VI), layer 5 (V), the cortical plate (CP), and the marginal zone (MZ). Immunofluorescence analysis of the neuronal transcription factors CUX1, TBR1, and CTIP2 at E16.5. Representative of 3 biological replicates. Scale bar = 100 μm. b) Morphology of E18.5 neurons after 1d in explant culture. Immunofluorescence staining for the pan-neuronal marker MAP2, tubulin beta 3 (TUBB3), and DAPI. Scale bar = 20 μm. c) Morphology of *Rad21^+/+^ Nex*^Cre^ and *Rad21*^lox/lox^ *Nex*^Cre^ cortical neurons in explant culture on rat glia (Kaech & Banker, 2006). Cultures were sparsely labeled with GFP to visualize individual cells and their processes, and stained for GAD67 to exclude GABAergic neurons. Dendritic traces of GFP^+^ neurons. Scale bar = 50 μm. d) Sholl analysis of *Rad21^+/+^ Nex*^Cre^ and *Rad21*^lox/lox^ *Nex*^Cre^ cortical neurons in explant cultures shown in c). Shown is the number of crossings, dendritic length, terminal points, and branch points per 10 μm. Three independent experiments, 32 *Rad21*^lox/lox^ *Nex*^Cre^ and 28 *Rad21^+/+^ Nex*^Cre^ neurons. * adj. *P* <0.05, ** adj. *P* <0.01, *** adj. *P* <0.001, **** adj. *P* <0.0001. Scale bar = 10 μm. e) Quantification of spines per 10 μm for *Rad21^+/+^ Nex*^Cre^ and *Rad21*^lox/lox^ *Nex*^Cre^ cortical neurons. Two independent experiments, 10 *Rad21*^lox/lox^ *Nex*^Cre^ and 10 *Rad21^+/+^ Nex*^Cre^ neuron). **** adj. *P* <0.0001. Scale bar = 10 μm.

Compared to freshly explanted E18.5 neurons (Fig. 3b), neurons acquired considerable morphological complexity after 14 days in explant culture (Fig. 3c). We sparsely labeled neurons with GFP to visualize processes of individual neurons (Fig. 3d) and used Sholl analysis (Sholl, 1953) to quantitate the number of axonal crossings, the length of dendrites, the number of terminal points, the number of branch points and the number of spines in GAD67-negative *Rad21*^+/+^ *Nex*^Cre^ and GAD67^-^negative *Rad21*^lox/lox^ *Nex*^Cre^ neurons. Cohesin-deficient neurons displayed reduced morphological complexity across scales, with reduced numbers of axonal branch and terminal points (Fig. 3c, d). In addition, *Rad21*^lox/lox^ *Nex*^Cre^ neurons showed reduced numbers of dendritic spines, the location of neuronal synapses (Fig. 3e). Taken together, these data show that the changes in neuronal gene expression that accompany cohesin deficiency have a tangible impact on neuronal morphology, and that cohesin is required for neuronal maturation.

### Activity-regulated gene expression is sensitive to acute depletion of cohesin

Neuronal maturation and ARG expression are closely connected: The expression of ARGs promotes neuronal maturation, morphological complexity, synapse formation, and connectivity. In turn, neuronal maturation, morphological complexity, synapse formation and connectivity facilitate ARG expression (Gallo et al., 2018; Kim et al., 2010; Malik et al., 2014; Greer et al., 2008; Tyssowski et al., 2018; Yap et al., 2018). To address whether the downregulation of ARGs observed in *Rad21*^lox/lox^ *Nex*^Cre^ neurons was cause or consequence of impaired maturation we examined gene expression changes 24 h after acute proteolytic degradation of RAD21-TEV depletion in post-mitotic neurons (Weiss et al., 2021). RNA-seq showed highly significant overlap in gene expression between acute degradation of RAD21-TEV and genetic cohesin depletion in *Rad21*^lox/lox^ *Nex*^Cre^ neurons (*P* < 2.22e-16, odds ratio = 8.24 for all deregulated genes, odds ratio = 23.81 for downregulated genes; Supplementary Fig. 5a). Differentially expressed genes were enriched for ontologies related to synapse function, cell adhesion, and neuronal/nervous system development, and showed expression changes that were correlated across cohesin depletion paradigms (Synapse: *P* < 2.22e−16, odds ratio = 11.26, *R_S_* = 0.62; Adhesion: *P* < 2.22e−16, odds ratio = 15.21, *R_S_* = 0.6; Neuronal and nervous system development: *P* < 2.22e−16, odds ratio = 9.92, *R_S_* =0.58; Supplementary Fig. 5b). Notably, ARGs were enriched among deregulated genes in response to acute RAD21-TEV cleavage (*P* <2.22e-16, Odds Ratio = 6.48), and were preferentially downregulated (Supplementary Fig. 5c). Thus, we conclude that activity-regulated genes and genes that facilitate neuronal maturation and homeostasis are directly affected by the acute depletion of cohesin, and not just as a result of impaired neuronal maturation.

### Activity-regulated gene classes differ with respect to their reliance on cohesin

The disruption of baseline ARG expression suggested that cohesin-deficient neurons may be unable to induce the same activity-dependent gene expression programme as wild-type neurons. To address this possibility, we performed RNA-seq of neuronal explant cultures treated either with tetrodotoxin + D-AP5 (TTX) alone to block neuronal signaling, or treated with TTX followed by 1 or 6h of sustained KCl exposure to induce neuronal depolarization. The fraction of constitutive genes deregulated in *Rad21*^lox/lox^ *Nex*^Cre^ versus control neurons remained similar across conditions (baseline, TTX, 1h and 6h KCl, Fig. 4a, top). By contrast, approximately 50% (154/305) of ARGs (Kim et al., 2010) were downregulated in *Rad21*^lox/lox^ *Nex*^Cre^ versus control neurons under baseline conditions (Supplementary Fig. 6a). Of these, 76 were induced to control expression levels by KCl in *Rad21*^lox/lox^ *Nex*^Cre^ neurons, while 29% failed to reach control levels and 16% were expressed at increased levels (Supplementary Fig. 6a). Multifactor analysis and hierarchical clustering (Supplementary Fig. 6b) showed that baseline ARG expression was more similar to TTX in *Rad21*^lox/lox^ *Nex*^Cre^ than in control neurons. This was confirmed by dendrogram distances (Supplementary Fig. 6c) and principal component analysis (Supplementary Fig. 6d), and statistically validated by the fraction of ARGs that changed expression between baseline and TTX conditions (49.5% in control neurons versus 28.5% in *Rad21*^lox/lox^ *Nex*^Cre^ neurons; *P* = 5.89e-11, Supplementary Fig. 6e). Taken together, these data show that ARG expression is reduced in *Rad21*^lox/lox^ *Nex*^Cre^ neurons under baseline conditions, but remains responsive to activation. Among neuronal ARG classes, IEGs and SRGs are known to differ in their 3D connectivity in that SRGs engage in longer-range chromatin contacts than IEGs in primary cortical neurons (Beagan et al., 2020). However, the functional consequences of this difference in 3D connectivity remain to be explored. We therefore asked whether IEGs and SRGs differ with respect to their reliance on cohesin. While IEGs were induced to at least wild-type levels in cohesin-deficient neurons by stimulation with KCl (Fig. 4a) or BDNF (Fig. 4b), a substantial fraction of SRGs remained downregulated in cohesin-deficient neurons across conditions (Fig. 4c). Expression of these SRGs was cohesin-dependent, as it was rescued by restoration of RAD21 levels following proteolytic cleavage of RAD21-TEV (Supplementary Fig. 7a). These data show that there are two classes of ARGs: (i) IEGs/SRGs that exhibit altered baseline expression but can fully regain expression in response to activation, and (ii) a subset of SRGs that remain deregulated in response to activation.

**Figure 4.**
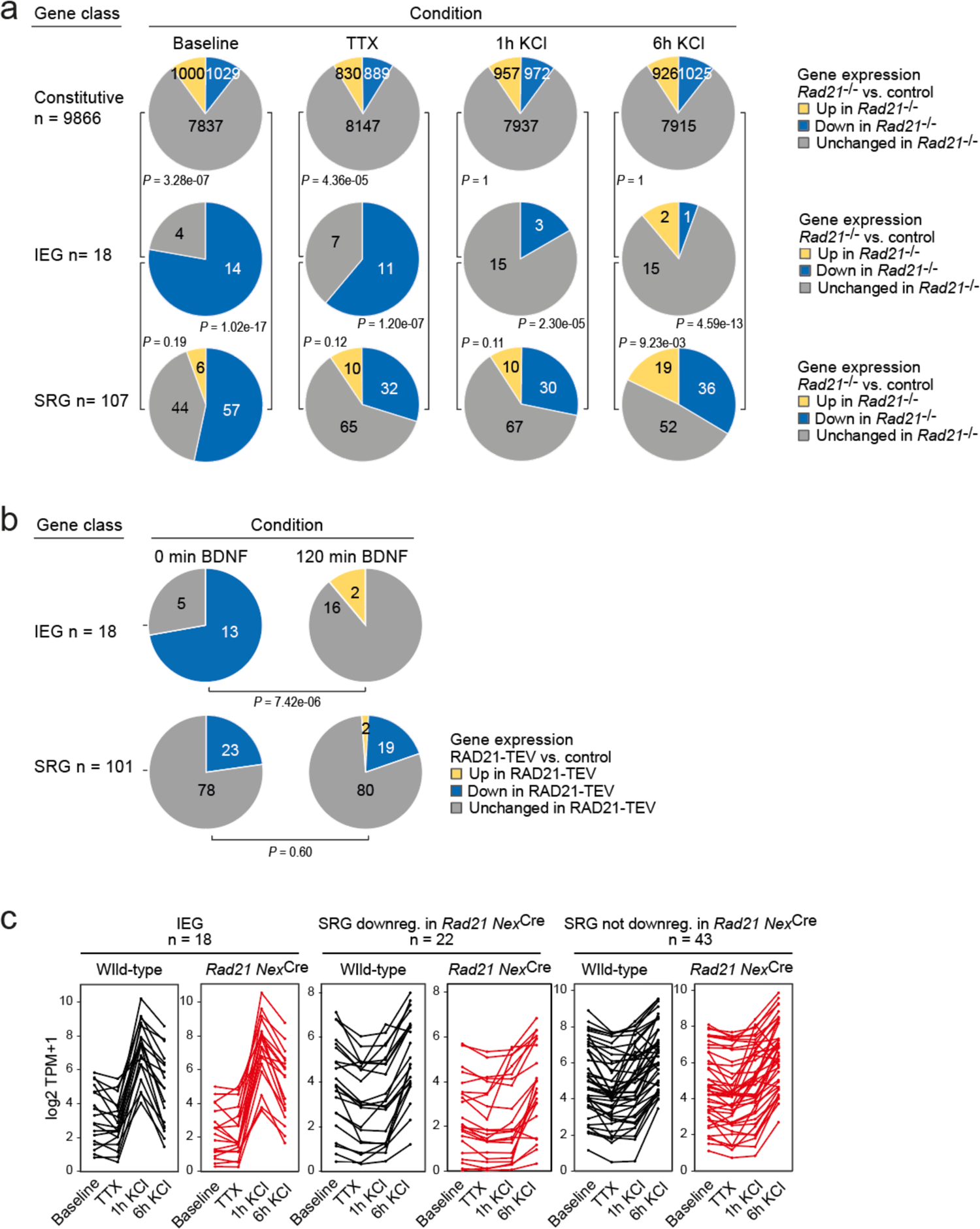
ARG classes differ in their reliance on cohesin. a) Pie charts show the expression of constitutive genes (top), IEGs (centre) and SRGs (bottom) in *Rad21* NexCre neurons under four different conditions: baseline, TTX and D-AP5 (TTX), and in response to KCl stimulation for 1h or 6h. Numbers of expressed constitutive genes, IEGs, and SRGs are given on the left. Note that KCl stimulation normalises the expression of most (10 out of 11) IEGs downregulated in TTX, but a fraction of SRGs remain downregulated. *P*-values test the prevalence of deregulated genes in each class under each condition, two-sided Fisher exact test. b) Pie charts show the expression of IEGs (top) and SRGs (bottom) in RAD21-TEV neurons under baseline conditions 24h after ERt2-TEV induction and in response to BDNF (120min). RAD21-TEV cleavage led to the downregulation (adj. P < 0.05) of 13 out of 18 expressed IEGs and of 23 out of 101 expressed SRGs. *P*-values test the prevalence of downregulated IEGs and SRGs with and without BDNF stimulation, two-tailed Fisher exact test. Note that BDNF stimulation reversed the downregulation of IEGs but not SRGs in cohesin-depleted neurons. c) Strip plots depict the expression of IEGs, SRGs that are downregulated in *Rad21* NexCre neurons compared to control across conditions (TTX and 6h KCl), and SRGs that are not downregulated in *Rad21* NexCre neurons compared to control across conditions.

### The dependence of neuronal gene expression scales with the genomic distance traversed by their chromatin loops

To explore features that may explain why a subset of ARG require cohesin for full expression we analysed ChIP-seq data for cohesin and CTCF binding to ARG promoters in wild-type neurons. ARG promoters are enriched for binding of CTCF (OR = 1.621, P = 0.022, two-tailed Fisher Exact Test), the cohesin subunit RAD21 (OR = 1.866, P = 0.005), and cohesin in the absence of CTCF (cohesin-non-CTCF, OR = 1.621, P = 0.022) compared to non-ARGs expressed in cortical neurons. Within the ARG gene set, however, there were no significant differences in CTCF, RAD21, or RAD21-non-CTCF ChIP-seq binding at the promoters of IEGs (which as a group remained fully inducible in cohesin-deficient neurons), the subset of SRGs that were downregulated across conditions in *Rad21* NexCre neurons (both TTX and 6h KCl, adj *P* < 0.05), and SRGs that remained fully inducible across conditions in *Rad21* NexCre neurons (both TTX and 6h KCl, adj *P* > 0.05, Supplementary Figure 7b). Therefore, while ARG promoters are enriched for CTCF, RAD21, and cohesin-non-CTCF binding, this binding is not predictive of which ARGs remain fully inducible, and which are downregulated in cohesin-deficient neurons.

To test whether cohesin-dependence of ARG regulation might instead be linked to the genomic range of chromatin loops formed by these genes we analysed high-resolution cortical neuron Hi-C data (Bonev et al., 2017). We found that the subset of SRGs that required cohesin for full expression in TTX and full induction by 6h KCl (adj *P* < 0.05) formed significantly longer Hi-C loops than IEGs and SRGs that were not deregulated across conditions (TTX and 6h KCl, adj *P* > 0.05) in *Rad21* NexCre neurons (Fig. 5a). Chromatin loops with CTCF binding at one or both loop anchors were also significantly longer for cohesin-dependent SRGs than for IEGs and cohesin-independent SRGs (Fig. 5a, red).

**Figure 5.**
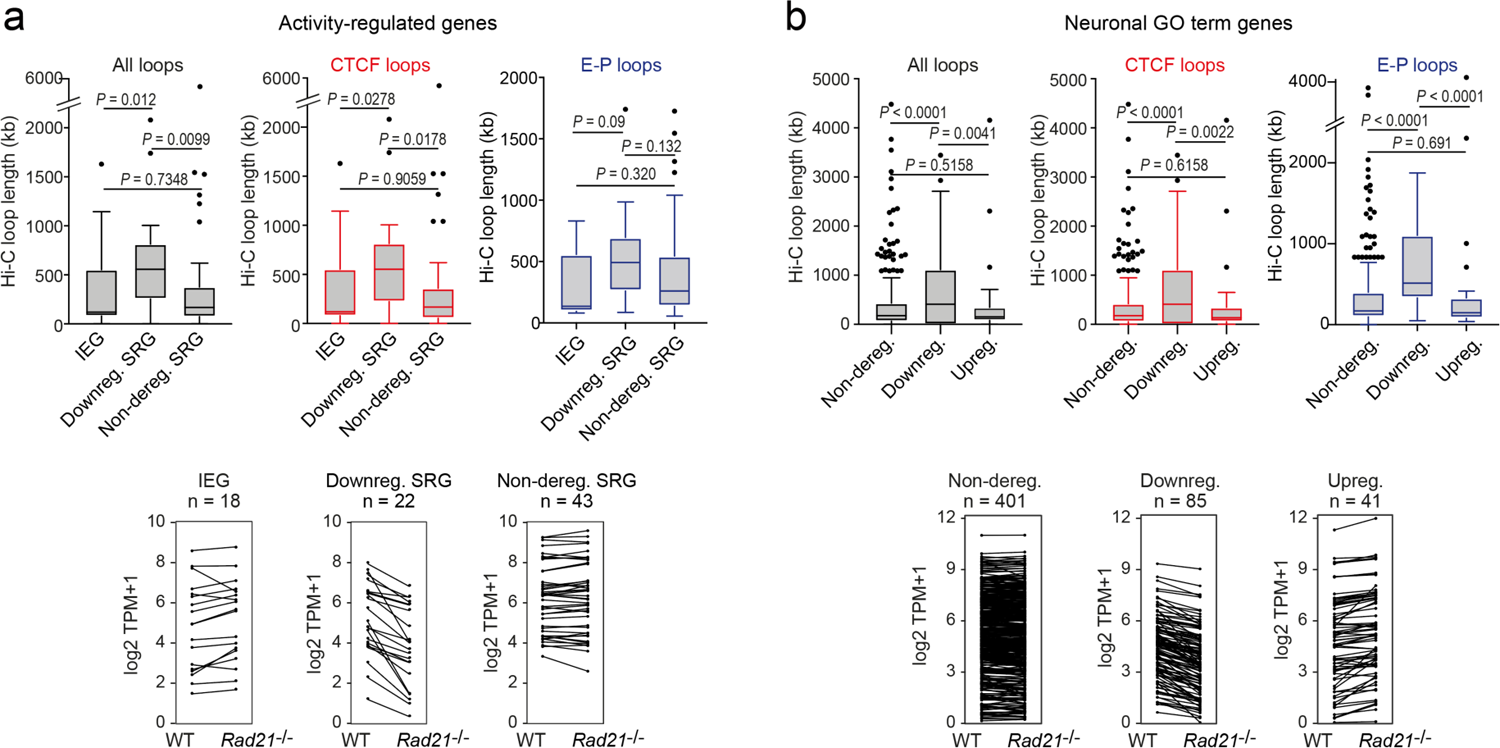
The genomic distance traversed by chromatin contacts formed by neuronal genes predicts whether or not cohesin is required for their full expression. a) The span of Hi-C loops (left), Hi-C loops with CTCF bound to at least one of the loop anchors (middle) and Hi-C loops between promoters and inducible enhancers (right) for IEGs (n=18) and SRGs downregulated in *Rad21 Nex*Cre versus control neurons in both resting (TTX) and activation conditions (6h KCl, adj P < 0.05 in both TTX and 6h KCl conditions, n = 22, ‘Downreg. SRG’), and SRGs not deregulated in either resting (TTX) or activation conditions (6h KCl, adj P > 0.05 in both TTX and 6h KCl conditions) in *Rad21 Nex*Cre relative to control neurons (adj. *P* > 0.05, n=43, ‘Non-dereg. SRG’). Box plots show the longest loop for each gene rather than average loop length, as Hi-C loop calling at 10kb resolution precludes detection of loops <40kb (Beagan et al., 2020). However, analysis of average loop length confirmed that downregulated genes form longer loops than non-deregulated genes among both ARGs and neuronal GO term genes (P = 0.0056 and P < 0.0001, respectively). Genes without loops are included except for analysis of enhancer loops. Box plots show the longest loop recorded for each gene.. Boxes show upper and lower quartiles and whiskers show 1.5 of the interquartile range. *P*-values were determined by non-parametric Kolmogorov-Smirnov test. Strip plots show IEG and SRG expression in control and *Rad21* NexCre neurons. Strip plots depict the expression of IEGs, downregulated SRGs and non-deregulated SRGs in wild-type and *Rad21* NexCre neurons. b) The span of Hi-C loops (left), Hi-C loops with CTCF bound to at least one of the loop anchors (middle) and Hi-C loops between promoters and constitutive or inducible enhancers (right) for genes in the neuronal GO terms synaptic transmission and glutamate receptor signaling. Gene expression in *Rad21 Nex*Cre versus control neurons was assessed in both resting and activation conditions: not deregulated in TTX or 6h KCl, n = 401, Downregulated in both TTX and 6h KCl, n = 85, Upregulated in both TTX and 6h KCl, n = 41). Genes do not form loops are included except for analysis of enhancer loops. Boxes show upper and lower quartiles and whiskers show 1.5 of the interquartile range. *P*-values were determined by non-parametric Kolmogorov-Smirnov test. Strip plots show the expression of the depicted GO term genes in control and *Rad21* NexCre neurons.

Chromatin loops that connect promoters with inducible enhancers also tended to span larger genomic distances at downregulated SRGs compared to IEGs or non-deregulated SRGs, even though due to the limited numbers of enhancer-promoter loops associated with each SRG class, these trends do not reach statistical significance (Fig. 5a, blue). Overall, the degree to which ARGs depend on cohesin for their correct expression correlates with the length of chromatin loops they form.

We next addressed whether cohesin/CTCF binding or chromatin loop length were important factors for the impact of cohesin on the expression of additional neuronal genes. We focused on the neuronal GO terms ‘synaptic transmission’ (GO:0007268) and ‘glutamate receptor signaling pathway’ (GO:0007215) because these gene ontologies were highly enriched among downregulated genes both in acute RAD21-TEV cleavage and genetic cohesin depletion, and remained enriched across conditions in *Rad21* NexCre neurons (GO:0007268: 87 of 519 genes downregulated at 6h KCl, adj. *P* < 0.05, *P*-value for enrichment = 2.99E-07; GO:0007215: 21 of 68 genes downregulated at 6h KCl, adj P<0.05, *P*-value for enrichment = 6.46E-06). The majority of synaptic transmission and glutamate receptor signaling genes are classified as constitutive (62.7% of expressed and 59.8% of downregulated GO:0007268 and GO:0007215 genes across conditions), rather than activity-regulated (1.8% of expressed and 1.2% of downregulated GO:0007268 and GO:0007215 genes cross conditions), thus complementing the analysis of ARGs. Of note, restoration of RAD21 after transient depletion in the RAD21-TEV system rescued the expression of 90% (70 of 77) downregulated synaptic transmission and glutamate receptor signaling genes (Supplementary Fig. 7c), confirming that their expression was indeed dependent on cohesin.

As described above for ARGs, the TSSs of neuronal GO term genes related to synaptic transmission and glutamate receptor signaling were enriched for binding of CTCF (OR = 1.409, P < 0.0001, two-tailed Fisher Exact Test), RAD21 (OR = 1.745, P < 0.0001), and cohesin-non-CTCF (OR = 1.585, P < 0.0001) compared to the remaining expressed genes in cortical neurons. However, and again as described above for ARGs, the binding of CTCF, RAD21, or cohesin-non-CTCF to the promoters of neuronal GO term genes did not predict which neuronal GO term genes remained expressed at wild-type levels, and which were downregulated in in *Rad21* NexCre neurons (Supplementary Fig. 7d).

Notably, however, synaptic transmission and glutamate receptor signaling genes that were downregulated in *Rad21^lox/lox^ Nex*^Cre^ neurons engaged in significantly longer-range chromatin loops than genes that were either not deregulated or upregulated (Fig. 5b, black, strip plots show the expression of the depicted GO term genes in control and *Rad21^lox/lox^ Nex*^Cre^ neurons). As with ARGs, this pattern extended to Hi-C loops with CTCF binding at one or both loop anchors (Fig. 5b, red). The subset of synaptic transmission and glutamate receptor signaling genes that were downregulated in *Rad21^lox/lox^ Nex*^Cre^ neurons formed significantly longer Hi-C loops connecting promoters and enhancers than genes in the same gene ontologies that were not deregulated (P<0.0001) or upregulated (P<0.0001) in *Rad21* NexCre neurons (Fig. 5b, blue). These data extend the relationship between chromatin loop length and cohesin-dependent expression from ARGs to constitutively expressed neuronal genes.

### Cohesin is not essential for short-range loops between inducible enhancers and promoters at the activity-regulated *Fos* and *Arc* loci

To examine the contribution of enhancer activation and enhancer-promoter contacts to inducible ARG expression in *Rad21*^lox/lox^ *Nex*^Cre^ neurons we focused on the immediate early response gene *Fos. Fos* expression in *Rad21*^lox/lox^ *Nex*^Cre^ neurons was reduced at baseline and in the presence of TTX, but *Fos* expression remained fully inducible when *Rad21*^lox/lox^ *Nex*^Cre^ neurons were stimulated with KCl (Fig. 6a) or with BDNF (Supplementary Fig. 8).

**Figure 6.**
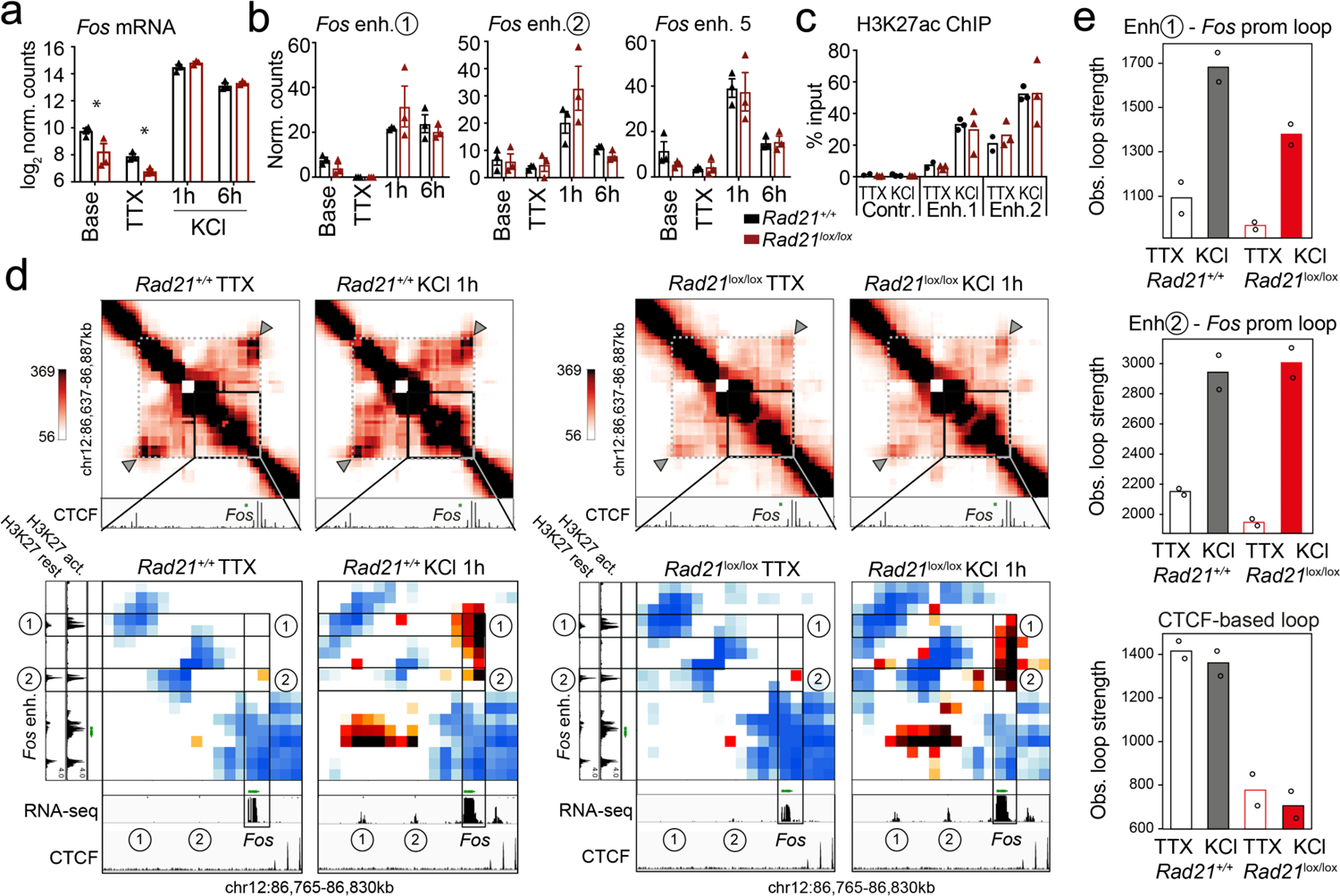
*Fos* enhancer-promoter contacts are robustly induced in cohesin-deficient neurons. a) Expression of the IEG *Fos* at baseline, after TTX/D-AP5 (TTX), and KCl-stimulation (left, mean log2-transformed counts from 3 biological replicates, * adj. *P* < 0.05). b) Enhancer transcripts in control and *Rad21*^lox/lox^ *Nex*^Cre^ neurons were quantified based on normalized RNA-seq reads within 1kb of the eRNA transcription start site. An intergenic region on chr11 was used as a negative control (71.177.622-71.177.792). c) H3K27ac ChIP normalized to H3 in control and *Rad21*^lox/lox^ *Nex*^Cre^ neurons at a control site, *Fos* enhancer 1 and *Fos* enhancer 2 after TTX/D-AP5 (TTX) or 1h KCl (KCl). d) Interaction score heatmaps of the 65 kb region immediately surrounding *Fos* obtained by 5C. Black frames highlight interactions between the *Fos* gene and upstream enhancers 1 and 2. CTCF ChIP-seq in cortical neurons (Bonev et al., 2017) is shown for orientation and H3K27ac ChIP-seq in inactive (TTX-treated) and activated neurons is shown to annotate enhancer regions (Beagan et al., 2020). RNA-seq in TTX-treated and 1h KCl-activated control and *Rad21*^lox/lox^ *Nex*^Cre^ neurons shows KCl-inducible transcription of *Fos* enhancers in wild-type and cohesin-deficient neurons. Two independent biological replicates are shown in Supplementary Fig. 10a. e) Quantification of 5C contacts between the *Fos* promoter and *Fos* enhancer 1 (top), the *Fos* promoter and *Fos* enhancer 2 (middle), and CTCF-marked boundaries of the sub-TAD containing *Fos* (bottom). Two replicates per genotype and condition.

The transcription of neuronal genes is controlled by neuronal enhancers (Rajarajan et al., 2016; Yamada et al., 2019; Sams et al., 2016; Kim et al., 2010; Malik et al., 2014; Schaukowitch et al., 2014; Beagan et al., 2020). Neuronal *Fos* enhancers in particular have been extensively characterised (Joo et al., 2016; Beagan et al., 2020), and interference with *Fos* enhancers precludes full induction of *Fos* gene expression (Joo et al., 2016). *Fos* enhancers 1, 2 and 5 are known to undergo activation-induced acetylation of H3K27 (H3K27ac) and active eRNA transcription in response to KCl stimulation (Joo et al., 2016). We found that activation-induced transcription of *Fos* enhancers did remain intact in *Rad21*^lox/lox^ *Nex*^Cre^ neurons (Fig. 6b) and activation-induced H3K27ac of *Fos* enhancers was also preserved (Fig. 6c).

Given that *Fos* can be fully induced, and *Fos* enhancers are activated in the absence of cohesin, we next set out to understand if *Fos* can form previously reported looping interactions with its activity-stimulated enhancers (Beagan et al., 2020). We conducted 5C to generate 10 kilobase resolution maps of chromatin loops around key IEGs and SRGs.

Consistent with previous data (Beagan et al., 2020), *Fos* enhancers 1 and 2, which are located ∼18 and ∼38.5 kb upstream of the *Fos* TSS, showed inducible chromatin contacts with the *Fos* promoter that formed rapidly in response to activation of wild-type neurons (Fig. 6d). Unexpectedly, *Rad21*^lox/lox^ *Nex*^Cre^ neurons retained the ability to robustly and dynamically induce loops between the *Fos* promoter and *Fos* enhancers 1 and 2 (Fig. 6d).

Quantification showed that inducible enhancer-promoter contacts at the *Fos* locus were of comparable strength in control and *Rad21*^lox/lox^ *Nex*^Cre^ neurons (Fig. 6e, top and center), while a structural CTCF-based loop surrounding the *Fos* locus was substantially weakened (Fig. 6e, bottom). Together these results reveal the surprising finding that the critical activity-stimulated IEG *Fos* can fully activate expression levels and form robust enhancer-promoter loops in the absence of cohesin.

We also examined the long-range regulatory landscape of the IEG *Arc*. The expression of the IEG *Arc* was reduced in *Rad21*^lox/lox^ *Nex*^Cre^ neurons at baseline, but, like *Fos, Arc* remained inducible by stimulation with KCl (Supplementary Fig. 9a, top) and BDNF (Supplementary Fig. 9a, bottom). Stimulation of wild-type neurons is known to trigger the formation of chromatin contacts between the *Arc* promoter and an *Arc*-associated activity-induced enhancer located ∼15kb downstream of the *Arc* TSS (Beagan et al., 2020; Supplementary Fig. 9b). As observed for *Fos*, *Arc* promoter-enhancer contacts were retained in *Rad21*^lox/lox^ *Nex*^Cre^ neurons (Supplementary Fig. 9b, c), while CTCF-based loops surrounding the *Arc* locus were weakened (Supplementary Fig. 9b, c). In contrast to control neurons, *Arc* promoter-enhancer contacts became at least partially independent of activation in *Rad21*^lox/lox^ *Nex*^Cre^ neurons (Supplementary Fig. 9b, c). Hence, cohesin is required for the correct baseline expression of ARGs, but largely dispensable for inducible transcription and for specific enhancer-promoter contacts at the IEGs *Fos* and *Arc*.

In contrast to IEGs *Fos* and *Arc*, the SRG *Bdnf* has at least eight promoters that initiate transcription of distinct mRNA transcripts, all of which contain the entire open reading frame for the BDNF protein (Aid et al., 2007). *Bdnf* promoter IV is specifically required for the neuronal activity-dependent component of *Bdnf* transcription in mouse cortical neurons (Hong et al., 2008). While overall *Bdnf* transcript levels were significantly reduced in *Rad21*^lox/lox^ *Nex*^Cre^ neurons only at baseline (log 2 FC = −1.16 adj *P* = 0.003, Supplementary Fig. 11a), *Bdnf* transcripts from the activity-dependent *Bdnf* promoter IV were specifically reduced in cohesin-deficient cortical neurons after 6h activation with KCl (log 2 FC = −1.26 adj P = 9.24e-05) as well as baseline conditions (log 2 FC = −1.39 adj *P* = 1.19e-05, Supplementary Fig. 11b). *Bdnf* promoter IV is located in the immediate vicinity of a strong CTCF peak in cortical neurons (Bonev et al., 2017; Supplementary Fig. 11c, bottom track). This CTCF peak forms the base of a constitutive loop between *Bdnf* promoter IV and an activity-induced enhancer located ∼2Mb upstream of the *Bdnf* gene (Supplementary Fig. 11c). The strength of this loop was substantially reduced in cohesin-deficient neurons (Supplementary Fig. 11c, quantification in Supplementary Fig. 11d). Hence, while the CTCF-based looping of *Bdnf* promoter IV to a distant inducible enhancer is cohesin-dependent, the activity-regulated expression of the IEGs *Fos* and *Arc* is linked to enhancer-promoter loops that span limited genomic distances (<40kb) and can form independently of cohesin.

## Discussion

Given that mutations in cohesin and CTCF cause intellectual disability in humans (Deardorff et al., 2018; Gregor et al., 2013; Rajarajan et al., 2016), the extent to which deficits in cohesin function alter neuronal gene expression remains a critical underexplored question. To define the role of cohesin in immature post-mitotic neurons we use experimental deletion of the cohesin subunit *Rad21* during a precise developmental window of terminal neuronal differentiation in vivo. We find impaired neuronal maturation and extensive downregulation of genes related to synaptic transmission, connectivity, neuronal development and signaling in *Rad21*^lox/lox^ *Nex*^Cre^ neurons. Such gene classes are central to neuronal identity, and their wide-spread downregulation is likely to contribute to the observed maturation defects of *Rad21*^lox/lox^ *Nex*^Cre^ neurons. Acute proteolysis of RAD21-TEV corroborated a prominent role for cohesin in the expression of genes that facilitate neuronal maturation, homeostasis, and activation.

We have recently shown that cohesin loss in macrophages results in severe disruption of anti-microbial gene expression programmes in response to macrophage activation (Cuartero et al., 2018). By contrast, cohesin loss only moderately affected genes constitutively expressed in uninduced and induced macrophages, supporting a model where cohesin-mediated loop extrusion is more important for the establishment of new gene expression than the maintenance of existing programmes (Cuartero et al., 2018). Here we extend this model to post-mitotic neurons in the murine brain. The two major ARG classes, IEGs and SRGs, were broadly downregulated in cohesin-deficient neurons at baseline. However, and in apparent contrast to inducible gene expression in macrophages, IEGs and a subset of SRGs remained fully inducible by KCl and BDNF stimulation in the absence of cohesin. Our results demonstrate that a subset of ARGs can undergo activity-dependent upregulation in the absence of cohesin. The reliance of ARGs on cohesin for full activity-induced expression was linked to the scale of chromatin interactions as quantified by the analysis of Hi-C loops. The subset of SRGs that exhibit defects in inducibility in the absence of cohesin is characterised by longer-range chromatin loops than either IEGs or SRGs that remain fully inducible in the absence of cohesin. The relationship between loop length and cohesin-dependence of neuronal gene expression extends to constitutively expressed cell type-specific neuronal genes. Neuronal genes related to synaptic transmission and glutamate receptor signaling that were downregulated in cohesin-deficient neurons also engaged in significantly longer chromatin loops than genes in the same GO terms that were not deregulated.

A subset of looping interactions made by ARGs and neuron-specific genes involve distal enhancers, suggesting that one role for cohesin in the expression of these genes may be to facilitate enhancer-promoter contacts. Our data indicate that specific enhancer-promoter loops at the key neuronal IEGs *Fos* and *Arc* can occur independently of cohesin in primary neurons. *Fos* enhancer-promoter loops remained responsive to environmental signals. Of note, *Fos* and *Arc* enhancer-promoter loops that were robust to cohesin depletion are relatively short-range (<40kb). By contrast, TAD-scale CTCF-based loops were substantial weakened, including a constitutive chromatin loop between *Bdnf* promoter IV and an activity-induced enhancer. In earlier studies we found that contacts between the *Lefty1* promoter and the +8kb enhancer, and between *Klf4* and enhancers at +53kb remained intact in acutely cohesin-depleted ES cells (Lavagnolli et al., 2015). Synthetic activation of a *Shh* enhancer ∼100kb upstream of the TSS supported transcriptional activation of cohesin, while activation of a +850kb enhancer did not (Kane et al., 2021). Analysis of engineered enhancer landscapes in K562 cells indicates graded distance effects: Enhancers at ≥ 100 kb and 47kb were highly and moderately dependent on cohesin, respectively, while loss of cohesin actually increased target gene transcription for enhancer distances ≤11kb (Rinzema et al., 2021). Finally, promoter capture Hi-C in cohesin-depleted HeLa cells indicates ranges of 10^4^-10^5^ bp for retained and 10^5^-10^6^ bp for lost interactions (Thiecke et al., 2020). While these studies suggest that cohesin-dependence of chromatin contacts relates to genomic distance, they fail to link this observation to physiologically relevant gene expression. The new data described here demonstrate scaling of cohesin-dependence with genomic distance in primary neurons, and, importantly, link this finding to critical genome functions, specifically the implementation of cell type-specific gene expression programmes during neuronal maturation and activation.

An open question concerns the mechanisms of enhancer-promoter contacts in the absence of cohesin. Current models of 3D genome folding posit competition between two forces, cohesin-mediated loop extrusion and condensate-driven compartmentalization (Rao et al., 2017; Nora et al. 2017; Schwarzer et al., 2017; Beagan & Phillips-Cremins, 2020). RNAP2, Mediator, the transactivation domains of sequence-specific transcription factors, and the C-terminal domain of the chromatin reader BRD4 are thought to support the formation of molecular condensates enriched for components of the transcriptional machinery (Sabari et al., 2018; Boija et al., 2018; Rowley et al., 2018; Hsieh et al., 2020). The activation of inducible ARG enhancers involves dynamic H3K27ac (Beagan et al., 2020), recruitment of RNA polymerases, and active transcription (Joo et al., 2016). Our data show that H3K27ac and transcription remain inducible at *Fos* enhancers in *Rad21*^lox/lox^ *Nex*^Cre^ neurons, which could potentially contribute to loop formation. At the *Arc* locus, the persistence of an enhancer-promoter loop in unstimulated cohesin-deficient neurons would be consistent with a role for cohesin-mediated loop extrusion in chromatin state mixing, and the separation of enhancer contacts (Rao et al., 2017). Notably, enhancer-promoter contacts resist the inhibition of BET proteins (Crump et al., 2021), selective degradation of BRD4 (Crump et al., 2021), Mediator (El Khattabi et al., 2019; Crump et al., 2021), RNA Polymerase II (Thiecke et al., 2020), or inhibition of transcription (El Khattabi et al., 2019). These observations suggest that numerous components of the transcriptional machinery may redundantly support associations between active genes and regulatory elements (Sabari et al. 2018; Boija et al. 2018). Nevertheless, phase separation-like forces provide an attractive hypothesis for the manner by which cohesin-independent enhancer-promoter loops may form.

In summary, our data demonstrate that the extent to which the establishment of activity-dependent neuronal gene expression programmes relies on cohesin-mediated loop extrusion depends at least in part on the genomic distances traversed by their long-range chromatin contacts.

## Acknowledgements

We thank Drs G. Little, M. Clements, and S. Parrinello (University College London) for advice and practical instruction, S. Di Giovanni and J. Merkenschlager (Rockefelller University) for comments on the manuscript, members of our labs for helpful discussions, L. Game for sequencing, and J. Elliott and B. Patel for cell sorting. This work was supported by the Medical Research Council UK, The Wellcome Trust (Investigator Award 099276/Z/12/Z to MM), EMBO (ALTF 1047-2012 TO LC), HFSP (LT00427/2013 to LC), the NIH National Institute of Mental Health (1R01-MH120269; 1DP1OD031253); JEPC), the NIH National Institute of Neural Disorders and Stroke (1R01-NS114226; JEPC), and a 4D Nucleome Common Fund grant (1U01DK127405; JEPC).

## Author contributions

LC, FDW, JAB, MSO, WG and KT did experiments, LC, FDW, JAB, MSO, Y-FW, TC, GD, KT, JEP-C and MM analysed data, LC, FDW, JAB, MAU, AGF, JEP-C and MM conceptualised the study, CW provided tools for image acquisition and analysis; LC, FDW, JAB, AGF, JEP-C and MM wrote the manuscript.

## Methods

### Mice

Mouse work was done under a UK Home Office project licence and according to the Animals (Scientific Procedures) Act. Mice carrying the floxed *Rad21* allele (*Rad21^lox^*, Seitan et al., 2011), in combination with the Cre recombinase in the Nex locus (*Nex^Cre^*, Goebbels et al., 2006) and where indicated *Rpl22(HA)^lox/lox^* RiboTag (Sanz et al., 2009) were on a mixed C57BL/129 background. For timed pregnancies the day of the vaginal plug was counted as day 0.5. Genotypes were determined by PCR as previously reported (Seitan et al., 2011; Sanz et al., 2009). *Rad21*^tev/tev^ mice have been described (Tachibana-Konwalski et al., 2010; Weiss et al., 2021).

### Neuronal cultures

For *Nex^Cre^* experiments, mouse cortices were dissected and dissociated from individual E17.5-E18.5 mouse embryos as described (Beaudoin et al., 2012) with minor modifications. Dissociated neurons were maintained in Neurobasal medium with B27 supplement (Invitrogen), 1 mM L-glutamine, and 100 U/mL penicillin/streptomycin for 10 days in vitro. Cells were plated at a density of 0.8 × 10^6^ cells per well on 6-well plates pre-coated overnight with 0.1 mg/ml poly-D-lysine (Millipore) and one third of the media in each well was replaced every 3 days. Cultures were treated with 5μM Cytosine β-D-arabinofuranoside (Ara-C, Sigma) from day 2-4. For immunofluorescence staining neurons were plated on 12 mm coverslips (VWR) coated with poly-D-lysine at a density of 0.1×10^6^ cells per coverslip. For cell-type-specific isolation of ribosome-associated mRNA, neurons from both cortices from each individual mouse embryo were seeded in a 10 cm dish.

For RAD21-TEV experiments, mouse cortices were dissected and dissociated on E14.5 – 15.5 as described (Weiss et al., 2021). Neurons were maintained in Neurobasal medium with B27 supplement (Invitrogen), 1mM L-glutamine, and 100U/ml penicillin/streptomycin. Cells were plated at a density of 1.25 × 10^5^/cm^2^ on 0.1mg/ml poly-D-lysine (Millipore) coated plates, and half the media was replaced every 3 days. Cultures were treated with 5μM Ara-C at day 5. For cleavage of RAD21-TEV, neurons were plated as described above and transduced at day 3 with lentivirus containing ERt2-TEV at a multiplicity of infection of 1. For ERt2-TEV dependent RAD21-TEV degradation, neurons were treated on culture day 10 with 500nM 4-hydroxytamoxifen (4-OHT) or vehicle (ethanol) for 24 hours.

For KCl depolarization experiments, neuronal cultures were pre-treated with 1μM tetrodotoxin (TTX, Tocris) and 100μM D-(-)-2-Amino-5-phosphonopentanoic acid (D-AP5, Tocris) overnight to reduce endogenous neuronal activity prior to stimulation. Neurons were membrane depolarized with 55 mM extracellular KCl by addition of prewarmed depolarization buffer (170 mM KCl, 2 mM CaCl2, 1 mM MgCl2, 10 mM HEPES pH7.5) to a proportion of 0.43 volumes per 1ml volume of neuronal culture medium in the well. For BDNF induction experiments, neuronal cultures were treated with BDNF (50ng/ml) for the indicated period of time at 10 days in vitro.

For Sholl analysis, dissociated cortical neurons were cultured as described (Greig et al., 2013). Astroglial monolayers were adhered to culture dishes and cortical neurons to coverslips, which were then suspended above the glia. Primary cultures of glial cells were prepared from newborn rat cortices. Four days before neuronal culture preparation, glial cells were seeded in 12-well plates at a density of 1×10^4^ cells per well and one day before, the medium from the glial feeder cultures was removed and changed to neuronal maintenance medium for preconditioning. Mouse cortices were dissected from E17.5/E18.5 mouse embryos and kept up to 24h in 2 mL of Hibernate™-E Medium (ThermoFisher) containing B27 supplement (Invitrogen) and 1 mM L-glutamine in the dark at 4°C. Embryos were genotyped and the cortices from the desired genotypes were used to prepare neuronal cultures as described before. Neurons were plated on 24-well plates containing poly-D-lysine precoated 12 mm coverslips (VWR) at a density of 0.1×10^6^ cells per well. Wax dots were applied to the coverslips, which served as ‘feet’ to suspend the coverslips above the glial feeder layer. Four hours after neuronal seeding, each coverslip containing the attached neurons was transfer upside down into a well of the 12-well dishes with the glial feeder.

Cultures were treated with 5μM Ara-C from day 2-4 and subsequently one third of the media was replaced every 3 days. For sparse neuronal GFP labelling, cortical neurons were transfected using 1μg of peGFP-N1 plasmid along with 2 μl per well of Lipofectamine 2000 (Invitrogen) after 12 days in culture. Cultures were maintained for 14 days *in vitro* before fixation.

### RNA extraction and RT-qPCR

RNA was extracted with QIAshredder and RNeasy minikit (Qiagen). Residual DNA was eliminated using DNA-free kit (Ambion) and reverse-transcribed using the SuperScript first-strand synthesis system (Invitrogen). RT-PCR was performed on a CFX96 Real-Time System (Bio-Rad) with SYBR Green Master Mix (Bio-Rad) as per the manufacturer’s protocol and normalized to *Ubc* and *Hprt* mRNA levels.

Relative level of the target sequence against the reference sequences was calculated using the ΔΔ cycle threshold method. RT–PCR primer sequences:

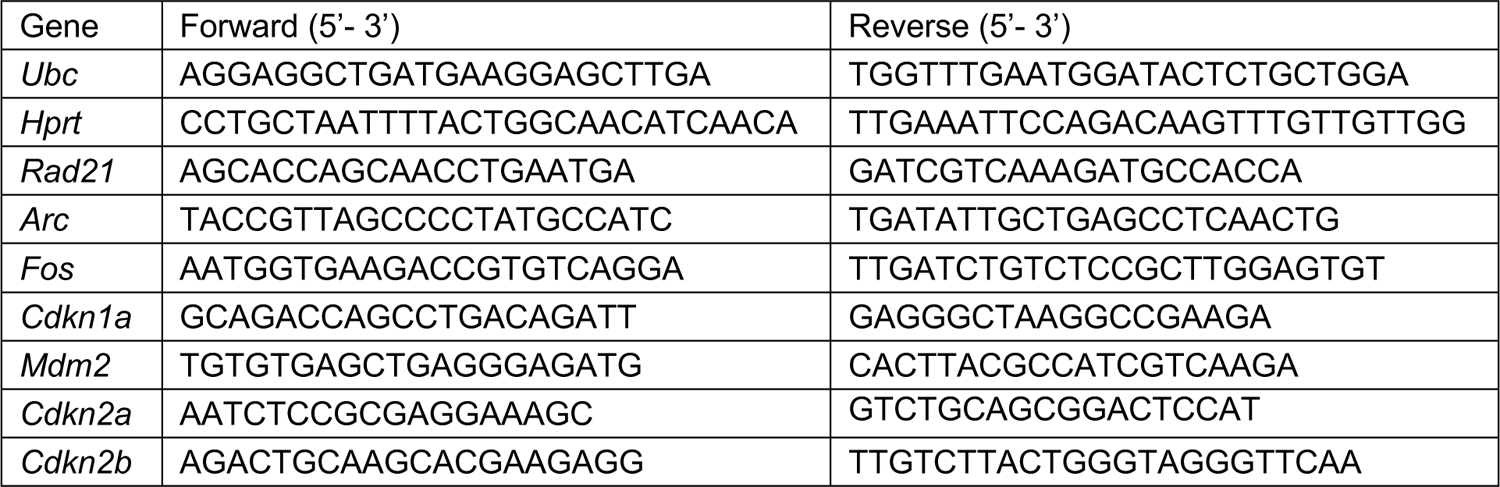

### Protein analysis

Whole cell extracts were prepared by resuspending cells in PBS with complete proteinase inhibitor (Roche, Cat#18970600), centrifugation, and resuspension in protein sample buffer (50mM Tris-HCl pH6.8, 1% SDS, 10% glycerol) followed by quantification using Qubit. Following quantification 0.001% Bromophenol blue and 5% beta-mercaptoethanol were added. Sodium dodecyl sulphate-polyacrylamide gel electrophoresis (SDS-PAGE) was carried out with the Bio-Rad minigel system. 20μg of protein sample and the benchmark pre-stained protein ladder (Biorad, #161-0374) were loaded on to a precast 10% polyacrylamide gel (Biorad, #456-1036). Resolved gels were blotted to a polyninylidene fluoride transfer membrane (Millipore, #IPVH00010) in transfer buffer (48mM Trizma base, 39mM glycine, 0.037% SDS and 20% methanol) using the trans-blot semi-dry electrophoretic transfer apparatus (BioRad). Membranes were incubated for 1 hour with fluorescent blocker (Millipore, HC-08) followed by primary antibody incubation diluted in blocker at an appropriate dilution for 2 hours or at room temperature or overnight at 4°C. Primary antibodies were rabbit polyclonal to RAD21 (1:1000; ab154769, Abcam), goat polyclonal to LAMIN B (1:10000; sc-6216; Santa Cruz Biotechnology), mouse monoclonal anti-myc tag (1:500, SC-40, Santa Cruz Biotechnology). Secondary antibodies were goat anti-rabbit IgG (H+L) Alexa Fluor 680 (1:10000; A-21109, ThermoFisher), goat anti-mouse IgG, Alexa Fluor 680 1:10,000), and donkey anti-goat IgG (H+L) Alexa Fluor 680 (1:10000; A-21084, ThermoFisher). Immobilon-FL PVDF membranes (Millipore) were imaged on an Odyssey instrument (LICOR).

### Cell-type-specific isolation of ribosome-associated mRNA

For polysome immunoprecipitation experiments, homogenates from 10 day cortical explant cultures were prepared as described (Sanz et al., 2009) with minor modifications. Cells were first washed two times on ice with 10 ml of PBS containing 100 μg/mL cycloheximide (Sigma). Cells were lysed in 50 mM Tris pH 7.5, 100 mM KCl, 12 mM MgCl2 (ThermoFisher), 1% IGEPAL CA-630 (Sigma), 1 mM DTT (Sigma), 200 U/mL RNasin (ThermoFisher), 1 mg/mL heparin (Sigma), 100 μg/mL cycloheximide, 1x Protease inhibitor (Sigma) and homogenization with a motor-driven grinder and pestle for about 2 minutes. Samples were then centrifuged at 10,000*g* for 10 min to create a postmitochondrial supernatant. For immunoprecipitations, 100 μL of Dynabeads™ Protein G (Invitrogen) were coupled directly to 10 μL of rabbit anti-HA antibody (Sigma, H6908). After polysome immunoprecipitation, total RNA was prepared using a RNeasy Plus Mini kit (Qiagen).

### RNAseq analysis

Total RNA was obtained in parallel from 10 day explant cultures of dissociated cortical neurons without stimulation (baseline); after overnight treatment with TTX and D-AP5 (TTX); and after overnight treatment with TTX and D-AP5 and depolarization with KCl for 1h (KCl1h) or 6h (KCl6h). RNA was extracted with QIAshredder and RNeasy mini kit (Qiagen). RNA-seq libraries were prepared from 600 ng of total RNA (RNA integrity number (RIN) >8.0) with TruSeq Stranded Total RNA Human/Mouse/Rat kit (Illumina). For polysome immunoprecipitation experiments, 300 ng of total RNA was used for library preparation (RIN>9.0). RNA from Rad21-TEV neurons was purified with a PicoPure RNA Isolation kit (Applied Biosystems KIT0204), and 200ng of total RNA was used to prepare libraries using the NEBNext® Ultra™ II Directional RNA Library Prep Kit for Illumina (polyA enrichment), following the manufacturer recommendations. Library quality and quantity were assessed on a Bioanalayser and Qubit respectively. Libraries were sequenced on an Illumina Hiseq2500 (v4 chemistry) and at least 40 million paired end 100bp reads per sample were generated per library and mapped against the mouse (mm9) genome. The quality of RNA-seq reads was checked by Fastqc (https://www.bioinformatics.babraham.ac.uk/projects/fastqc/) and aligned to mouse genome mm9 using Tophat version 2.0.11 (Kim et al., 2013) with parameters “--library-type=fr-firststrand”. Gene coordinates from Ensembl version 67 were used as gene model for alignment. Quality metrics for the RNA-Seq alignment were computed using Picard tools verion 1.90 (https://broadinstitute.github.io/picard/). Genome wide coverage for each sample was generated using bedtools genomeCoverageBed and converted to bigwig files using bedGraphToBigWig application from UCSC Genome Browser. Bigwig files were visualised using IGV. After alignment, number of reads on the genes were summarised using HTSeq-count (version 0.5.4; Anders et al., 2015). All downstream analysis was carried out in R (version 3.4.0). Differentially expressed genes between condition were determined using DESEq2 (Love et al., 2014). P-values calculated by DESeq2 were subjected to multiple testing correction using Benjamini-Hochberg method. Adjusted p-value of 0.05 was used to select the differentially expressed genes. Principal Component Analysis (PCA) and hierarchical clustering of samples were done on the normalised read counts (rlog) computed using DESeq2. KCl-inducible genes were defined as genes in *Rad21^+/+^* neurons with adj*P* <0.05 and log2 fold change ≥1 in KCl1h versus (vs) TTX or KCl6h vs TTX. As reference we used previously defined activity dependent genes (Kim et al., 2010). Constitutive genes were defined as expressed genes in wild-type neurons with adj. *P* ≥0.05 in KCl1h vs TTX and KCl6h vs TTX.

GO terms enriched among differentially expressed genes were identified using goseq R package (Young et al., 2010) using all expressed genes in each comparison as background. GSEA was performed as described (Subramanian et al., 2005) using GSEA Desktop v3.0 (http://www.broadinstitute.org/gsea). ‘Wald statistics’ from DESeq2 differential expression analysis were used to rank the genes for GSEA. The gene set collections C2 (curated; KEGG 186 gene sets) and C5 (GO ontologies; 5917 gene sets) were obtained from Molecular Signature Database (MSigDB version 6.1; Broad Institute, http://www.broadinstitute.org/gsea/msigdb). Neuron-specific genes were identified using Neutools (Gao et al., 2018).

### Chromatin immunoprecipitation

ChIP was performed as described with minor modifications. Briefly, cells were cross-linked for 10 minutes at room temperature with rotation using 1% formaldehyde in cross-linking buffer (0.1 M NaCl, 1 mM EDTA, 0.5 mM EGTA and 25 mM HEPES-KOH, pH 8.0). The reaction was quenched using 125 mM glycine for 5 minutes with rotation and the cells were washed three times using ice-cold PBS containing complete protease inhibitor cocktail tablets (Roche). Cells were resuspended in lysis buffer (1% SDS, 50 mM Tris-HCl pH 8.1, 10 mM EDTA pH 8) with EDTA-free protease inhibitor cocktail and incubated for 30 minutes on ice. Cell lysates were then sonicated 20 times at 4°C (Bioruptor Plus, Diagenode, 30/30 cycles) and centrifuged at 14000 rpm for 1 minute at 4°C to remove cellular debris. 10% of total volume was taken as input. Input samples were reverse crosslinked overnight at 65°C, then incubated for 1 h with 9 mM EDTA pH 8, 3.6 mM Tris-HCl pH 6.8 and 36 μg/mL proteinase K at 45°C. Input chromatin purification was performed using phenol-chloroform at 4°C. Total chromatin was pre-cleared for 1h at 4°C with rotation using protein A sepharose beads (P9424, Merck). 3 µg of anti-histone H3 (Abcam, ab1791) and 5 µg of anti-H3K27Ac (Active Motif, 39133) were added overnight at 4°C with rotation. 100 μL of protein A sepharose beads were added to each IP for at least 4 h before being washed with low salt buffer (150 mM NaCl, 2 mM Tris pH 8.1, 0.1% SDS, 1% Triton X-100 and 2 mM EDTA pH 8), high salt buffer (500 mM NaCl, 20 mM Tris pH 8.1, 0.1% SDS, 1% Triton X-100 and 2 mM EDTA pH 8), LiCl salt buffer (0.25 mM LiCl, 10 mM Tris pH 8.1, 1% NP-40, 1% sodium deoxycholate and 1 mM EDTA pH 8) and Tris-EDTA buffer (10 mM Tris pH 8.1 and 1 mM EDTA pH 8). Chelex-100 (Bio-Rad, catalog number #1421253) was added to the samples, which were then boiled and incubated for 1 h at 55°C with 36 μg/mL proteinase K and boiled once again. Samples were centrifuged at 12000 rpm for 1 minute and the beads washed once with nuclease-free water. The following PCR primers were used: Control region chr11: 71.177.622-71.177.792: forward, 5’-CATTCCAGGGCAACTCCACT-3′, reverse, 5’-CAGGGGCTCCTGTACTACCT-3′; *Fos* enhancer 1 forward, 5′-TCCGGTAAGGGCATTGTAAG-3′, reverse, 5′-CAAAGCCAGACCCTCATGTT-3′; *Fos* enhancer forward, 5′-TGCAGCTCTGCTCCTACTGA-3′, reverse, 5′-GAGGAGCAAGACTCCCACAG-3′

### 3C Template Generation

Neuronal cultures were fixed in 1% formaldehyde for 10 minutes (room temp) via the addition (1:10 vol/vol) of the following fixation solution: 50 mM Hepes-KOH (pH 7.5), 100 mM NaCl, 1 mM EDTA, 0.5 mM EGTA, 11% Formaldehyde. Fixation was quenched via the addition of 2.5 M glycine (1:20 vol/vol) and scraped into pellets. Each pellet was washed once with cold PBS, flash frozen, and stored at −80°C. For each condition, *in situ* 3C was performed on 2 replicates of 4-5 million cells as described (Beagan et al., 2020). Briefly, cells were thawed on ice and resuspended (gently) in 250 μL of lysis buffer (10 mM Tris-HCl pH 8.0, 10 mM NaCl, 0.2% Igepal CA630) with 50 μL protease inhibitors (Sigma P8340). Cell suspension was incubated on ice for 15 minutes and pelleted. Pelleted nuclei were washed once in lysis buffer (resuspension and spin), then resuspended and incubated in 50 μL of 0.5% SDS at 62°C for 10 min. SDS was inactivated via the addition of 145 μL H_2_O, 25 uL 10% Triton X-100, and incubation at 37°C for 15 min.

Subsequently, chromatin was digested overnight at 37°C with the addition of 25 μL 10X NEBuffer2 and 100U (5 μL) of HindIII (NEB, R0104S), followed by 20 min incubation at 62°C to inactivate the HindIII. Chromatin was re-ligated via the addition of 100 μL 10% Triton X-100, 120 μL NEB T4 DNA Ligation buffer (NEB B0202S), 12 μL 10 mg/mL BSA, 718 μL H_2_O, and 2000 U (5 μL) of T4 DNA Ligase (NEB M0202S) and incubation at 16°C for 2 hours. Following ligation nuclei were pelleted, resuspended in 300 μL of 10 mM Tris-HCl (pH 8.0), 0.5 M NaCl, 1% SDS, plus 25 μL of 20 mg/mL proteinase K (NEB P8107), and incubated at 65°C for 4 hours at which point an additional 25 μL of proteinase K was added and incubated overnight. 3C templates were isolated next day via RNaseA treatment, phenol-chloroform extraction, ethanol precipitation, and Amicon filtration (Millipore MFC5030BKS). Template size distribution and quantity were assessed with a 0.8% agarose gel.

### 5C Library Preparation

5C primers which allow for the query of genome folding at ultra-high resolution but on a reduced subset of the genome were designed according to the double-alternating design scheme (Kim et al., 2018; Beagan et al., 2020) using My5C primer design (Lajoie et al., 2009; http://my5c.umassmed.edu/my5Cprimers/5C.php) with universal “Emulsion” primer tails. Regions were designed to capture TAD structures immediately surrounding the genes of interest in published mouse cortex HiC data (Bonev et al., 2017). 5C reactions were carried out as previously described (Beagan et al., 2020). 600 ng (∼200,000 genome copies) of 3C template for each replicate was mixed with 1 fmole of each 5C primer and 0.9 ug of salmon sperm DNA in 1x NEB4 buffer, denatured at 95°C for 5 min, then incubated at 55°C for 16 hours. Primers which had then annealed in adjacent positions were ligated through the addition of 10 U (20 μL) Taq ligase (NEB M0208L) and incubation at 55°C for 1 hour then 75°C for 10 min. Successfully ligated primer-primer pairs were amplified using primers designed to the universal tails (FOR = CCTCTC TATGGGCAGTCGGTGAT, REV = CTGCCCCGGGTTCCTCATTCTCT) across 30 PCR cycles using Phusion High-Fidelity Polymerase. Presence of a single PCR product at 100 bp was confirmed via agarose gel, then residual DNA <100 bp was removed through AmpureXP bead cleanup at a ratio of 2:1 beads:DNA (vol/vol). 100 ng of the resulting 5C product was prepared for sequencing on the Illumina NextSeq 500 using the NEBNext Ultra DNA Library Prep Kit (NEB E7370) following the manufacturer’s instructions with the following parameter selections: during size selection, 70 μL of AMPure beads was added at the first step and 25 at the second step; linkered fragments were amplified using 8 PCR cycles. A single band at 220 bp in each final library was confirmed using an Agilent DNA 1000 Bioanalyzer chip, and library concentration was determined using the KAPA Illumina Library Quantification Kit (#KK4835). Finally, libraries were evenly pooled and sequenced on the Illumina NextSeq 500 using 37 bp paired-end reads to read depths of between 11 and 30 million reads per replicate.

### 5C Interaction Analysis

5C analysis steps were performed as described (Gilgenast et al., 2019; Beagan et al., 2020; Fernandez et al., 2020). Briefly, paired-end reads were aligned to the 5C primer pseudo-genome using Bowtie, allowing only reads with one unique alignment to pass filtering. Only reads for which one paired end mapped to a forward/left-forward primer and the other end mapped to a reverse/left-reverse primer were tallied as true counts. Primer-primer pairs with outlier count totals, resulting primarily from PCR bias, were identified as those with a count at least 8-fold higher (100-fold for the lower-quality Arc region) than the median count of the 5×5 subset of the counts matrix centered at the primer-primer pair in question; outlier counts were removed.

Primer-primer pair counts were then converted to fragment-fragment interaction counts by averaging the primer-primer counts that mapped to each fragment-fragment pair (max of 2 if both a forward/left-forward and a reverse/left-reverse primer were able to be designed to both fragments and were not trimmed during outlier removal). We then divided our 5C regions into adjacent 4 kb bins and computed the relative interaction frequency of two bins (i,j) by summing the counts of all fragment-fragment interactions for which the coordinates of one of the constituent fragments overlapped (at least partially) a 12 kb window surrounding the center of the 4 kb i^th^ bin and the other constituent fragment overlapped the 12 kb window surrounding the center if the j^th^ bin. Binned count matrices were then matrix balanced using the ICE algorithm^69,71^ and quantile normalized across all 8 replicates (2 per condition) within each experimental set (neuronal activation and cohesin-rescue experimental datasets were quantile normalized separately) as previously described (Beagan et al., 2020), at which point we considered each entry (i,j) to represent the Observed Interaction Frequency of the 4 kb bins i and j. Finally, the background contact domain ‘expected’ signal was calculated using the donut background mode (Su et al., 2017) and used to normalize the relative interaction frequency data for the background interaction frequency present at each bin-bin pair. The resulting background-normalized interaction frequency (“observed over expected”) counts were fit with a logistic distribution from which p-values were computed for each bin-bin pair and converted into ‘Background-corrected Interaction Scores’ (interaction score = −10*log_2_(p-value)), which have previously shown to be informatively comparable across replicates and conditions (Beagan et al., 2020).

### Identification of neuronal enhancers

H3K27Ac ChIPSeq and corresponding input datasets (Malik et al., 2014) were from NCBI GEO using accession GSE60192. The sra files were converted to fastq using sratoolkit and aligned to mouse genome mm9 using bowtie version 0.12.8 with default parameters (Langmead et al., 2009). Sequencing reads aligned to multiple positions in the genome were discarded. Duplicate reads were identified using Picard tools v 1.90 and removed from the downstream analysis. H3K27Ac peaks were identified using macs2 with “--broad” parameters (Zhang et al., 2008). Gene coordinates were obtained from Ensembl using “biomaRt” R package (Durninck et al., 2009) and enhancers were assigned to nearest genes using “nearest” function from GenomicRanges R package (Lawrence et al., 2013). Enhancers at Hi-C loop anchors were defined as reported previously (Beagan et al., 2020) and are provided there as Table S11.

### Hi-C Loop Span Analysis

Loops were called on mouse cortical neuron HiC data (Bonev et al., 2017) as described (Beagan et al., 2020, loop calls are provided as Table S16 in Beagan et al., 2020). Cortical neuron CTCF ChIP-seq (Bonev et al., 2017) reads were downloaded from GEO accessions GSM2533876 and GSM2533877, merged, and aligned to the mm9 genome using Bowtie v0.12.7. Reads with more than two possible alignments were removed (-m2 flag utilized). Peaks were identified using MACS2 version 2.1.1.20160309 with a P value cutoff parameter of 10^−4^. Loops were classified as ‘CTCF loops’ if a CTCF peak fell within at least one anchor of the loop. Loops were classified as enhancer-promoter loops if the TSS of a gene of interest fell within one loop anchor and the other anchor of the same loop contained an activity-induced enhancer (for ARGs) or a constitutive or activity-induced enhancer (for neuronal genes related to synaptic transmission and glutamate receptor signaling; Table S11 in Beagan et al., 2020). The span of loops with an anchor which contained the TSS of a gene of interest was quantified by calculating the genomic distance between the midpoint of the loop’s two anchors.

### Immunocytochemistry

Neurons plated on coverslips were fixed with warmed to 37^0^C PBS containing 4% paraformaldehyde and 4% sucrose for 10 min at room temperature. Neurons were then permeabilized with 0.3% Triton X-100 for 10 min and treated with blocking solution (10% normal goat serum, 0.1% Triton X-100 in PBS) for one hour. Primary and secondary antibodies were diluted in 0.1% Triton X-100, 2% normal goat serum in PBS. Appropriate primary antibodies were incubated with samples for two hours at room temperature or overnight at 4°C. Primary antibodies used were specific to RAD21 (1:500; rabbit polyclonal ab154769, Abcam), MAP2 (1:5000; chicken polyclonal ab611203, Abcam), GAD67 (1:500; mouse monoclonal MAB5406, Millipore), HA (1:1000; mouse monoclonal MMS-101R, Covance), GFAP (1:500; rabbit polyclonal Z0334, Dako) or IBA1 (1:250; rabbit polyclonal 019-19741, Wako). Secondary antibodies were incubated for 1 h at room temperature. Goat anti-rabbit IgG (H+L) Alexa Fluor 647 (A-21244, ThermoFisher), Goat anti-Rabbit IgG (H+L) Alexa Fluor 568 (A-11011, ThermoFisher), goat anti-mouse IgG (H+L) Alexa Fluor 488 (A-11001, ThermoFisher), goat anti-chicken IgY (H+L) Alexa Fluor 568 (ab175711, Abcam) conjugates were used at a 1:500 dilution. Cells were mounted in Vectashield medium containing DAPI (Vector Labs).

Embryonic brains were fixed for 4 hours in 4% paraformaldehyde in PBS, washed in PBS, transferred to 15% sucrose in PBS for cryopreservation, embedded in OCT and stored at −80°C until use. Coronal sections of 10 μm were cut with a Leica cryostat and mounted on glass slides. The sections were washed two times for ten minutes in PBS, blocked with 0.3% TritonX-100, 5% normal goat serum in PBS, at room temperature and incubated overnight with the primary antibody solution. Sections were then washed three times for ten minutes each in PBS and were incubated for one hour in the dark with the secondary antibody solution. The primary antibodies were specific to RAD21 (1:500; rabbit polyclonal ab154769, Abcam), anti-gamma-H2AX (1:3000; rabbit polyclonal A300-081A, Bethyl Laboratories), Cleaved Caspase-3 (Asp175) (1:400; rabbit polyclonal 9661, Cell signalling), TBR1 (1:1000; rabbit polyclonal ab31940, Abcam), CTIP2 (1:500; rat monoclonal [25B6] ab18465, Abcam), CUX-1 (1:400; rabbit polyclonal sc-13024, Santa Cruz) and anti phospho-histone H3 (Ser10) Alexa Fluor 647 conjugate (1:50; rabbit polyclonal 9716, Cell signalling). The secondary antibodies used are described in the previous section. Sections were mounted with Vectashield medium containing DAPI.

### Confocal Image analysis and quantification

For quantification analysis of RAD21 negative neurons and inhibitory neurons (GAD67+) images (1024 × 1024 pixels) were acquired using TCS SP5 confocal microscope (Leica Microsystems), using a HCX PL APO CS 40x/1.25 lens at zoom factor 1 (373 nm/pixel). Images were acquired with identical settings for laser power, detector gain, and amplifier offset, with pinhole diameters set for 1 airy unit. DAPI-identified nuclei that colocalized with the GAD67 signal, or without RAD21 signal were counted as inhibitory or RAD21 negative neurons, respectively; and were quantified using a processing pipeline developed in CellProfiler (version 2.2, Broad Institute, Harvard, Cambridge, MA, USA, www.cellprofiler.org). For astrocytes (GFAP+) and microglia (IBA1+) quantification in dissociated cortical neuronal cultures, the entire coverslips were imaged using a IX70 Olympus microscope equipped with a 4x 0.1 NA Plan-Neofluar lens (1.60 μm/pixel) and the number of astrocytes and microglia were counted. The analysis and quantification of different cell types were done for three different experiments, each experiment containing at least two different samples of each genotype.

For Sholl analysis of dissociated cortical neurons, samples were imaged using a TCS SP8 confocal microscope, a HC PL APO CS2 40x/1.30 lens at zoom factor 0.75 with a resolution of 2048 x 2048 pixel (189 nm/pixel, 0.5 μm/stack). Images were acquired with identical settings for laser power, detector gain, and amplifier offset, with pinhole diameters set for 1 airy unit. Approximately 10 neurons were imaged per sample and at least two different samples per genotype in each experiment; each experiment was performed three times.

GFP+ neurons were traced using the FilamentTracer package in Imaris software (Bitplane AG). Statistical significance was assessed using a repeat measures ANOVA with a Bonferroni Post Test (Prism-GraphPad Software).

## Supplementary Figures

**Supplementary Figure 1.**
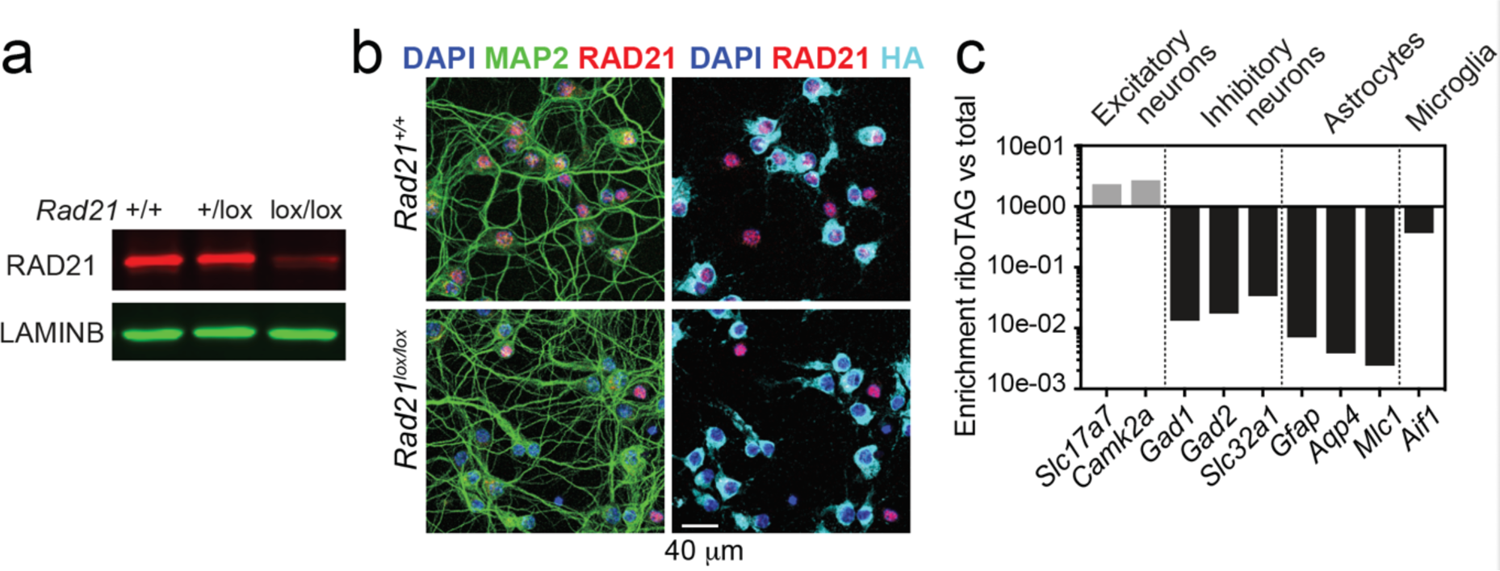
*Rad21* Nex^Cre^ RiboTag validation. a) RAD21 and LAMIN B protein expression in *Rad21^+/+^ Nex*^Cre^ and *Rad21*^lox/lox^ *Nex*^Cre^ cortical explant cultures was analysed by fluorescent immunoblots. One representative blot of six is shown here, the quantification of all 6 blots is shown in Fig. 1d. b) *Nex*^Cre^-dependent Rpl22-HA (RiboTag) expression is restricted to RAD21-negative cells in *Rad21*^lox/lox^ *Nex*^Cre^ neurons. Immunofluorescence staining for RAD21, the pan-neuronal marker MAP2 and HA (RiboTag) in explant culture. DAPI marks nuclei. Scale bar = 40 μm. c) *Nex*^Cre^ RiboTag captures excitatory neuron-specific transcripts such as *Slc17a7* and *Camk2a* and depletes cell type-specific transcripts expressed in inhibitory neurons (*Gad1, Gad2, Slc32a1*), astrocytes (*Gfap, Aqp4, Mlc1*), and microglia (*Aif*). Transcript enrichment (or depletion) was calculated using the normalized counts from *Nex*^Cre^ RiboTag versus standard RNA-seq in *Rad21^+/+^ Nex*^Cre^ neurons.

**Supplementary Figure 2.**
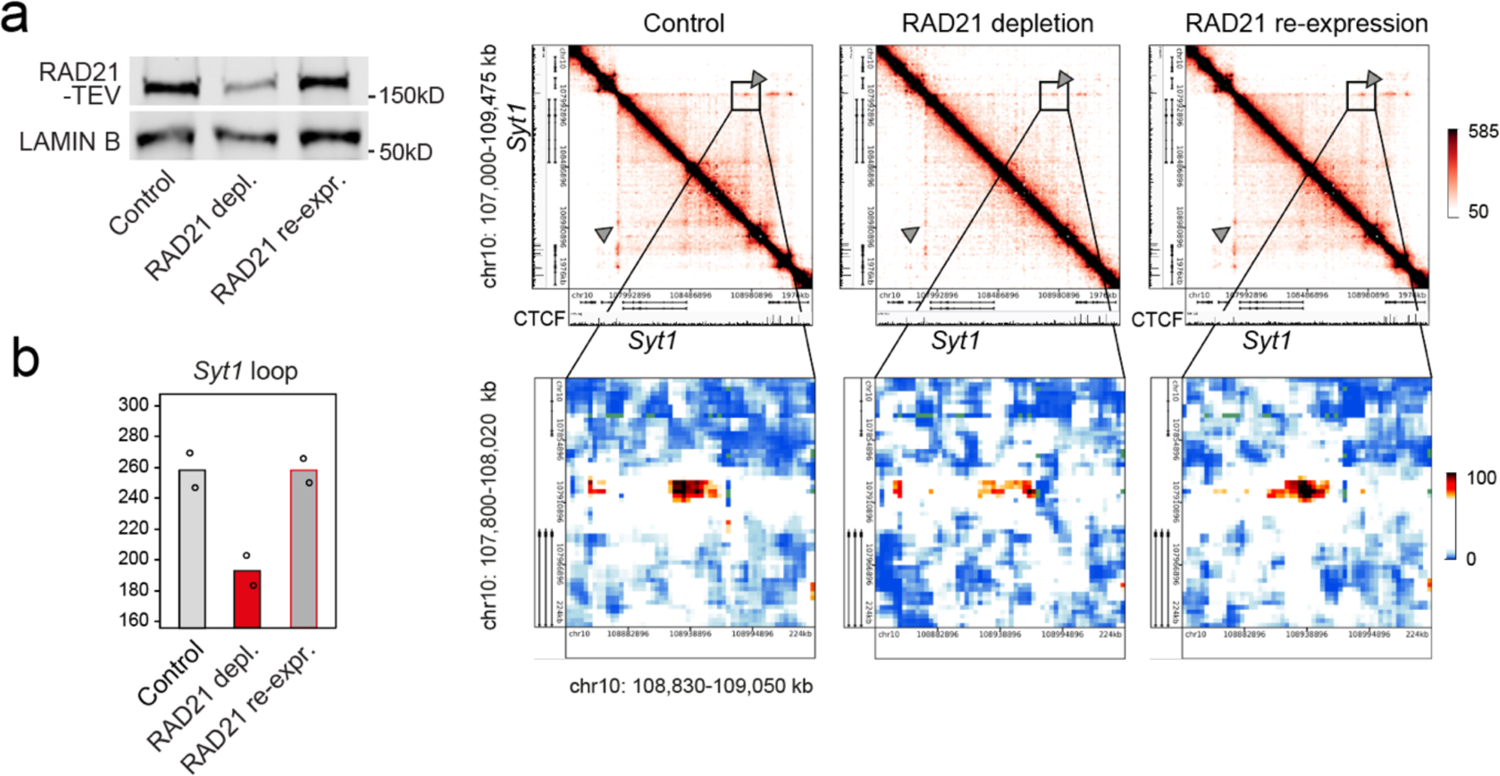
Restoration of cohesin rescues chromatin loops. a) Transient RAD21 depletion by Dox-inducible TEV expression and recovery after Dox washout (Weiss et al., 2021). RAD21-TEV western blots (left) and heat maps of chromatin contacts at the cohesin-dependent neuronal genes *Syt1* in RAD21-TEV neurons (right). b) Quantification of chromatin loops at *Syt1* in control neurons, cohesin-depleted neurons 24h after Dox-dependent TEV induction, and after re-expression of RAD21.

**Supplementary Figure 3.**
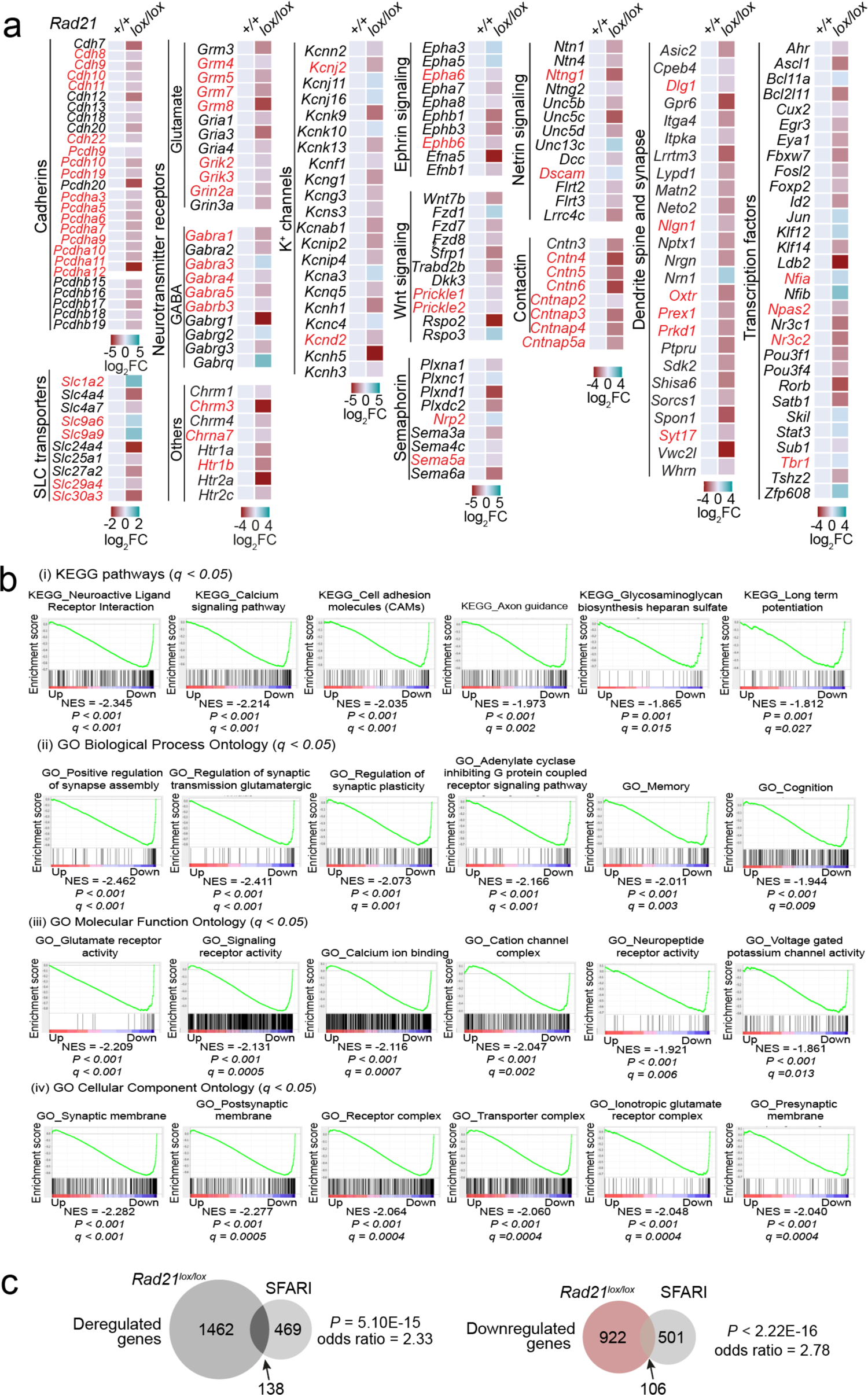
Gene expression in *Rad21*^lox/lox^ *Nex*^Cre^ neurons. a) Examples of deregulated genes in *Rad21*^lox/lox^ *Nex*^Cre^ neurons. Genes associated with autism spectrum disorders are highlighted in red. d) GSEA for downregulated genes in *Nex^Cre/+^ Rad21^lox/lox^* neurons using gene sets derived from (i) KEGG pathway database, (ii) GO Biological Process Ontology, (iii) GO Molecular Function Ontology, (iv) GO Cellular Component Ontology in the Molecular Signatures Database (MSigDB). c) Overlap between human genes associated with autism spectrum disorders from the SFARI database and differentially expressed genes (left), downregulated genes (middle) and upregulated genes (right) in *Rad21*^lox/lox^ *Nex*^Cre^ cortical neurons.

**Supplementary Figure 4.**
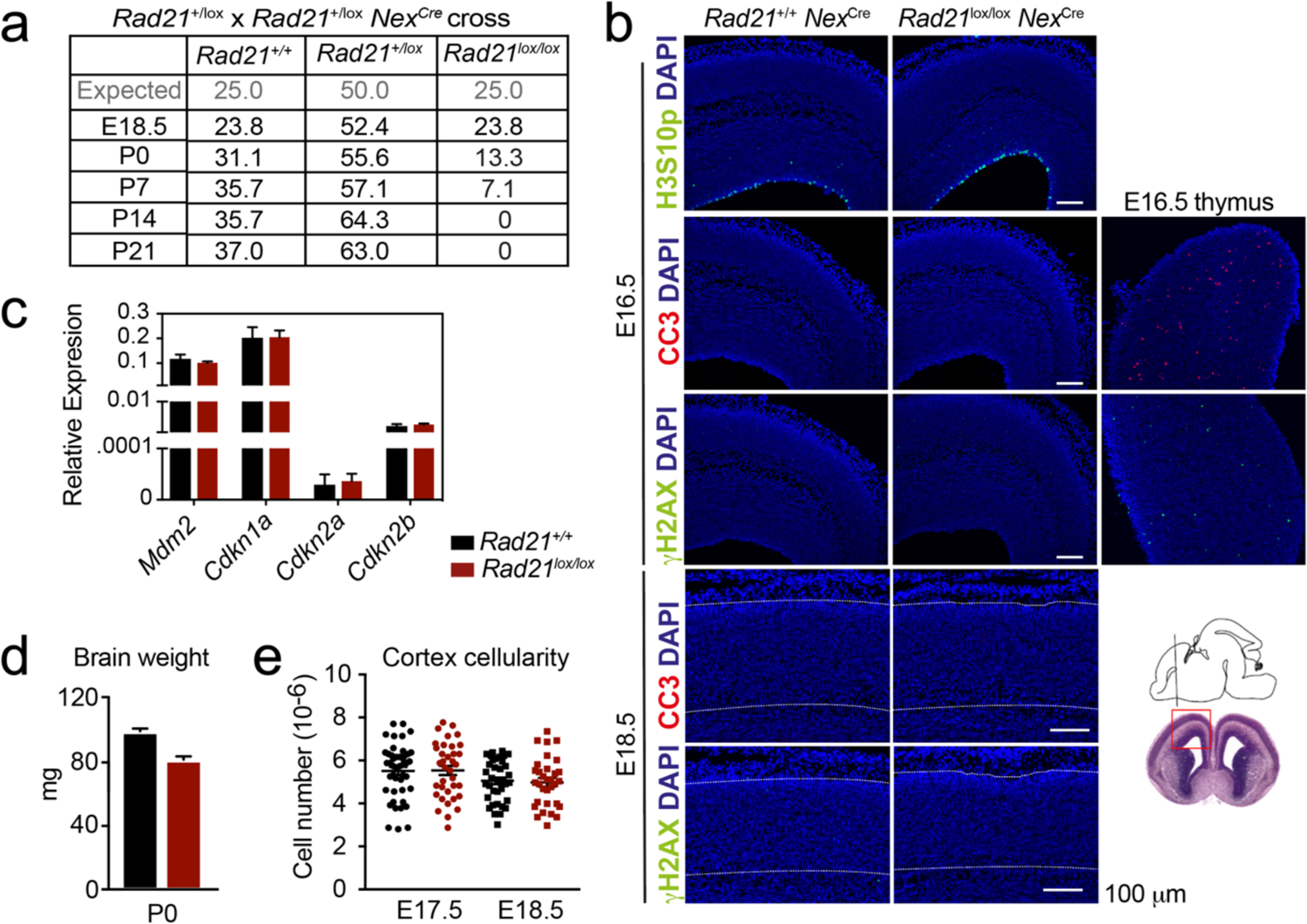
Impact of cohesin loss in immature post-mitotic neurons *in vivo*. a) Expected Mendelian ratios and observed percentages of live *Rad21*^+/+^ *Nex*^Cre^*, Rad21*^lox/+^ *Nex*^Cre^, *Rad21*^lox/lox^ *Nex*^Cre^ mice at the indicated developmental stages, n = 217. b) Immunofluorescence analysis shows that neither the mitotic marker phosphorylated Serine 10 on histone H3 (H3S10p) nor the apoptosis marker activated caspase 3 (CC3) or the DNA damage marker γH2AX in E16.5 (top) and E18.5 *Rad21*^lox/lox^ *Nex*^Cre^ (bottom, white lines demarcate the cortex). Wild type E16.5 thymi are shown as positive controls for CC3 and γH2AX. Two biological replicates. Scale bar = 100 μm. Photomicrographs of coronal sbrain sections at gestational age E16 modified from the Atlas of the prenatal moue brain (Paxinos et al., 2020) are shown for orientation. c) Quantitative RT-PCR analysis of gene expression in *Rad21^+/+^ Nex*^Cre^ and *Rad21*^lox/lox^ *Nex*^Cre^ E17.5/18.5 cortical explant cultures 10 d after plating. *Hprt* and *Ubc* were used for normalization. Mean ± SEM of 3 cultures per genotype. d) Brain weights of *Rad21*^+/+^ *Nex*^Cre^ and *Rad21*^lox/+^ *Nex*^Cre^, *Rad21*^lox/lox^ *Nex*^Cre^ mice at birth (P0). Mean ± SEM of between 3 and 6 mice per genotype. e) Embryonic cortices from wild-type and *Rad21*^lox/lox^ *Nex*^Cre^ mice were dissected at E17.5 and E18.5 and dissociated. Cortical cell numbers were determined by counting in Neubauer chambers. Each symbol denotes an independent experiment. Mean ± SEM are also shown.

**Supplementary Figure 5.**
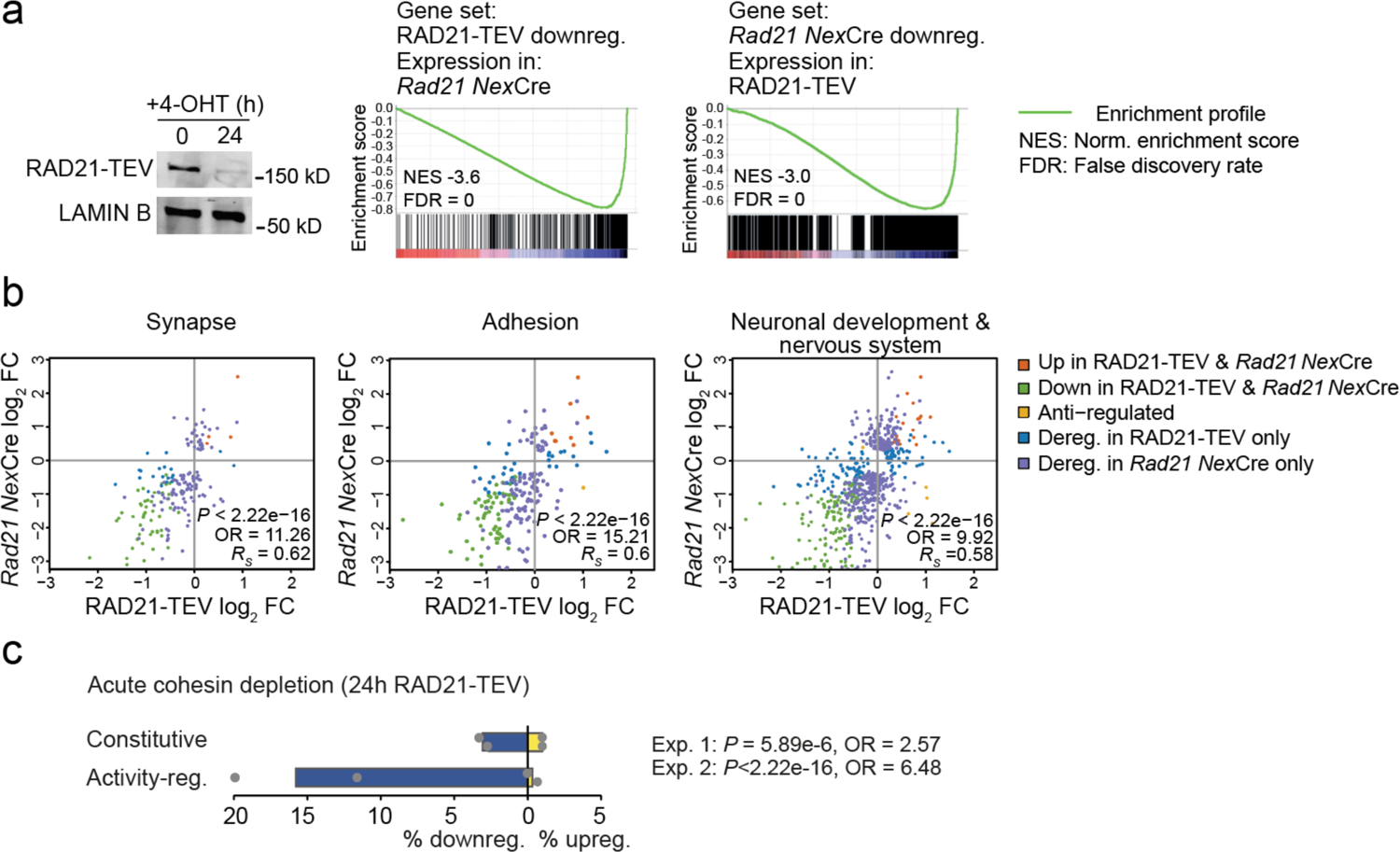
Activity-regulated gene expression is sensitive to acute cohesin depletion. a) Western blot documenting acute RAD21 depletion by 4-OHT-inducible RAD-TEV cleavage (left). GSEA of the gene set downregulated (DEseq2, adj. *P* < 0.05) in RAD21-TEV neurons in *Rad21*^lox/lox^ *Nex*^Cre^ neurons (centre). GSEA of genes downregulated in *Rad21*^lox/lox^ *Nex*^Cre^ neurons (DEseq2, adj. *P* < 0.05) in RAD21-TEV neurons (right). NES: normalised enrichment score. FDR: false discovery rate. b) Scatter plots of gene expression within aggregate GO terms, comparing RAD21-TEV with *Rad21*^lox/lox^ *Nex*^Cre^ neurons. Genes that were found deregulated in at least one of the genotypes are shown. *P*-values and odds ratios refer to the probability of finding the observed patterns of co-regulation by chance. *R_S_*: Spearman’s rank coefficient. c) Deregulation of constitutive and activity-regulated genes 24h after acute cohesin depletion by inducible proteolytic cleavage of RAD21-TEV; adj. *P* <0.05 based on DEseq2 analysis of 3 RNA-seq replicates per experiment. Blue indicates downregulation and yellow indicates upregulation in RAD21-TEV versus wild-type. Two independent experiments are shown (Weiss et al., 2021 and Supplementary Data 4).

**Supplementary Figure 6.**
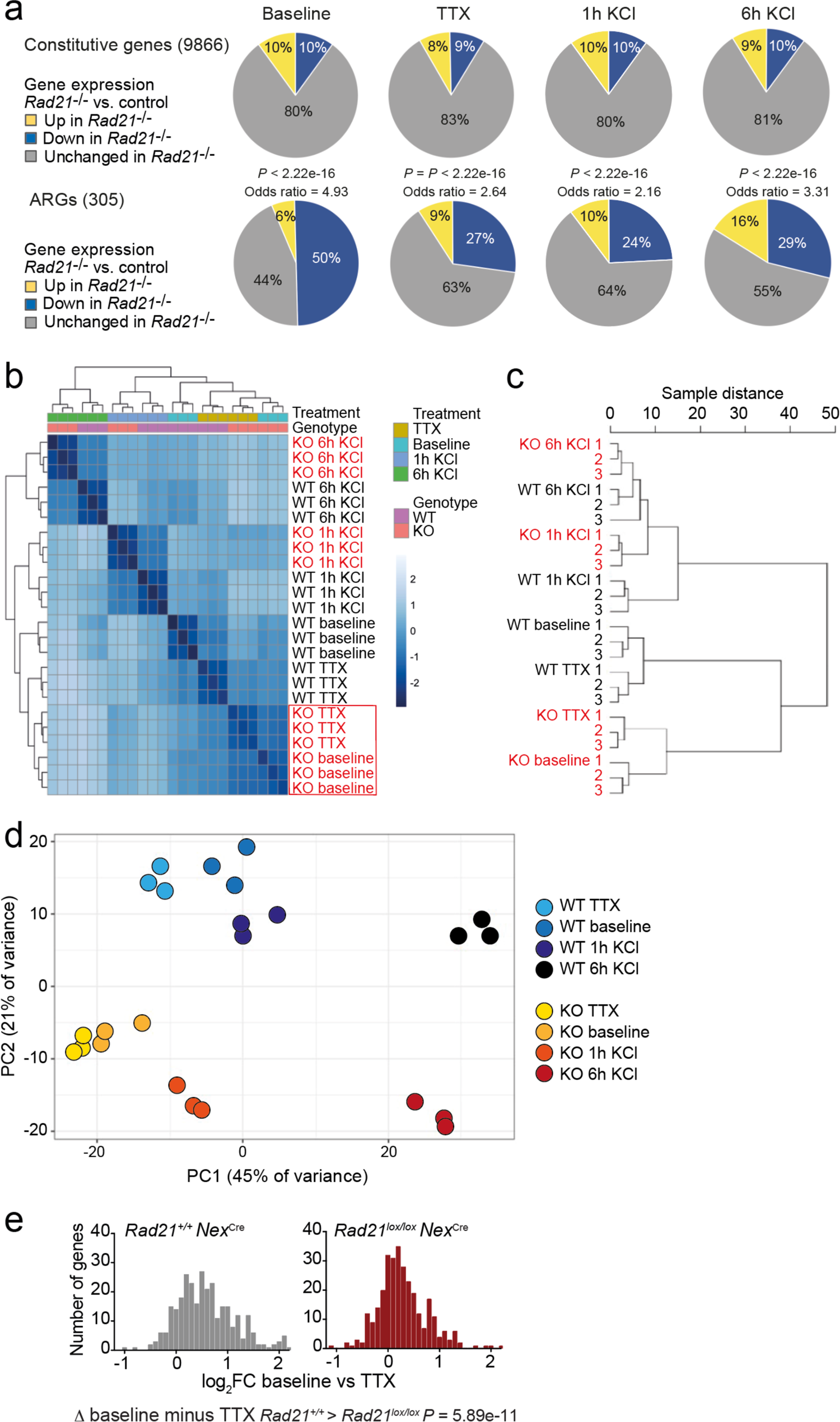
Gene expression and genotype interaction analysis. a) The expression of constitutive and previously defined activity-regulated genes (Kim et al., 2010) was determined by RNA-seq in explant cultures of *Rad21*^lox/lox^ *Nex*^Cre^ neurons at baseline, in the presence of TTX and D-AP5 (TTX), and after stimulation with KCl for 1h and 6h. Odds ratios show the enrichment of ARGs. *P*-values for deviation of odds ratios from 1 were determined by 2 tailed Fisher’s Exact test. b) Heatmap of sample distances for genotype interaction analysis of *Rad21*^+/+^ and *Rad21*^lox/lox^ *Nex*^Cre^ neurons across conditions. Note the clustering of baseline and TTX conditions for *Rad21*^lox/lox^ *Nex*^Cre^ neurons (red box) c) Principal component analysis of *Rad21*^+/+^ and *Rad21*^lox/lox^ *Nex*^Cre^ neurons across conditions (all genes). PC1 (45% of variance) separates activation conditions and PC2 separates genotype (21% of variance). d) Sample distance dendrograms indicate that ARG expression in *Rad21*^lox/lox^ *Nex*^Cre^ neurons shows little difference between spontaneous activity at baseline and TTX. e) ARG expression in explant cultures of *Rad21*^+/+^ and *Rad21*^lox/lox^ *Nex*^Cre^ neurons under baseline conditions that allow for cell-cell communication versus TTX/D-AP5 (TTX). The difference in ARG expression between baseline and TTX was greater for wild-type than for *Rad21*^lox/lox^ *Nex*^Cre^ neurons (*P* = 5.89e-11, Kolmogorov-Smirnov test).

**Supplementary Figure 7.**
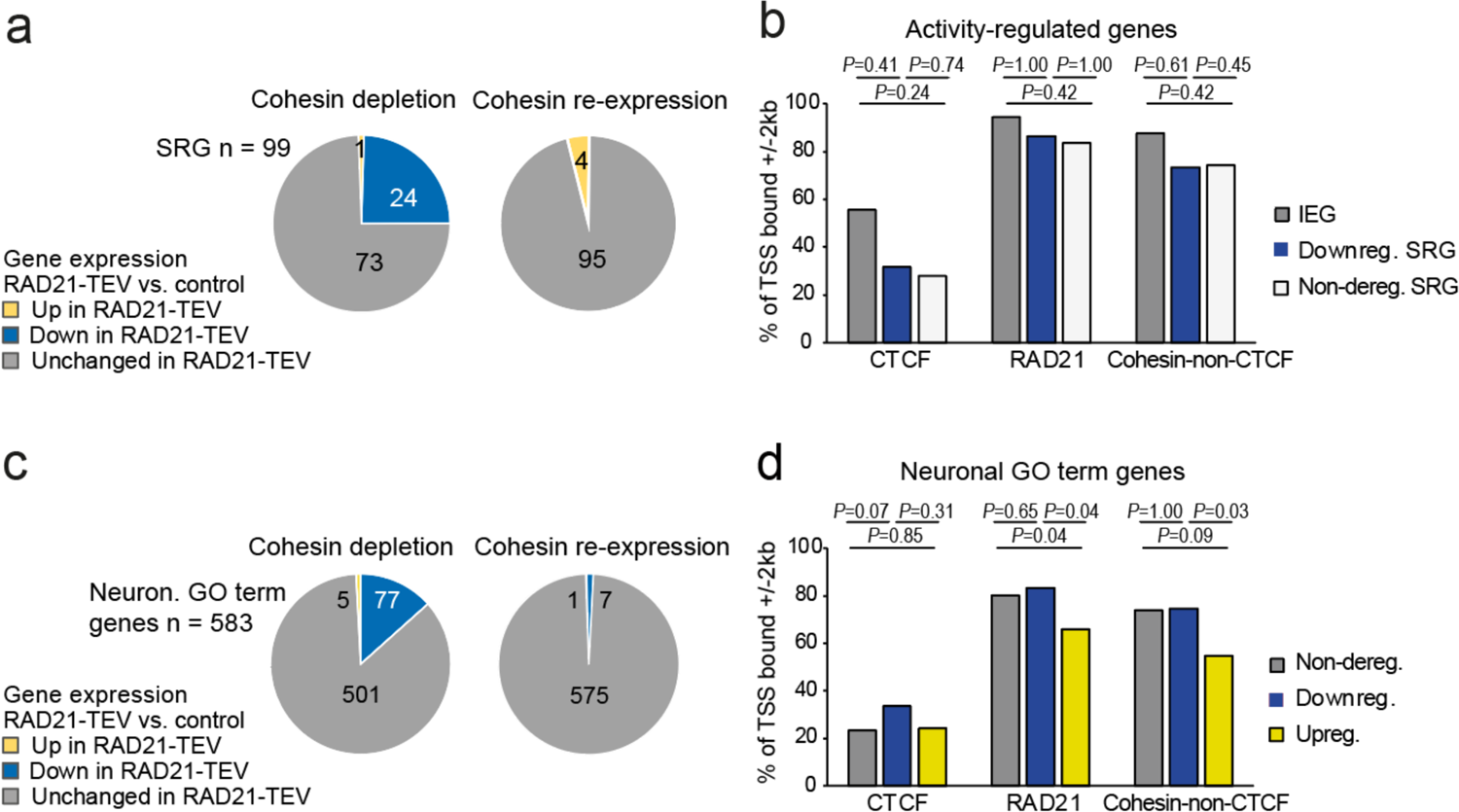
Restoration of cohesin rescues the expression of neuronal genes. a) Transient cohesin depletion and re-expression as in Supplementary Fig. 2. Pie charts show the expression of SRGs in RAD21-TEV relative to control neurons. 24 out of 97 SRGs expressed in RAD21-TEV neurons were downregulated 24h after Dox-dependent TEV induction (adj. *P* < 0.05). The downregulation of SRGs was reversible upon Dox washout and restoration of RAD21 expression. b) Presence of CTCF (Bonev et al., 2017), RAD21 (Fujita et al., 2017), or cohesin-non-CTCF binding at gene promoters (TSS ± 2kb) at IEGs, SRGs that are downregulated across conditions (TTX, 6h KCl), and SRGs that not deregulated across conditions in *Rad21 Nex*Cre versus control neurons (left). *P*-values: two-tailed Fisher exact test. c) Transient cohesin depletion and re-expression as in Supplementary Fig. 2. Pie charts show the expression of neuronal genes related to synaptic transmission (GO:0007268) and glutamate receptor signaling pathway (GO:0007215). 77 out of 583 expressed genes in these GO terms (Neur. GO term genes) were downregulated 24h after Dox-dependent TEV induction (adj. *P* < 0.05). The downregulation of 76 of 77 neuronal GO term genes was reversible upon Dox washout and restoration of RAD21 expression, an additional 6 genes were downregulated after Dox washout but not at 24h of TEV induction. d) Presence of CTCF (Bonev et al., 2017), RAD21 (Fujita et al., 2017), or cohesin-non-CTCF binding at gene promoters (TSS ± 2kb) at neuronal genes related to ‘synaptic transmission’ (GO:0007268) and ‘glutamate receptor signaling pathway’ (GO:0007215) that are not deregulated across conditions, downregulated across conditions, or upregulated across conditions (TTX, 6h KCl) in *Rad21 Nex*Cre versus control neurons (right). *P*-values: two-tailed Fisher exact test.

**Supplementary Figure 8.**
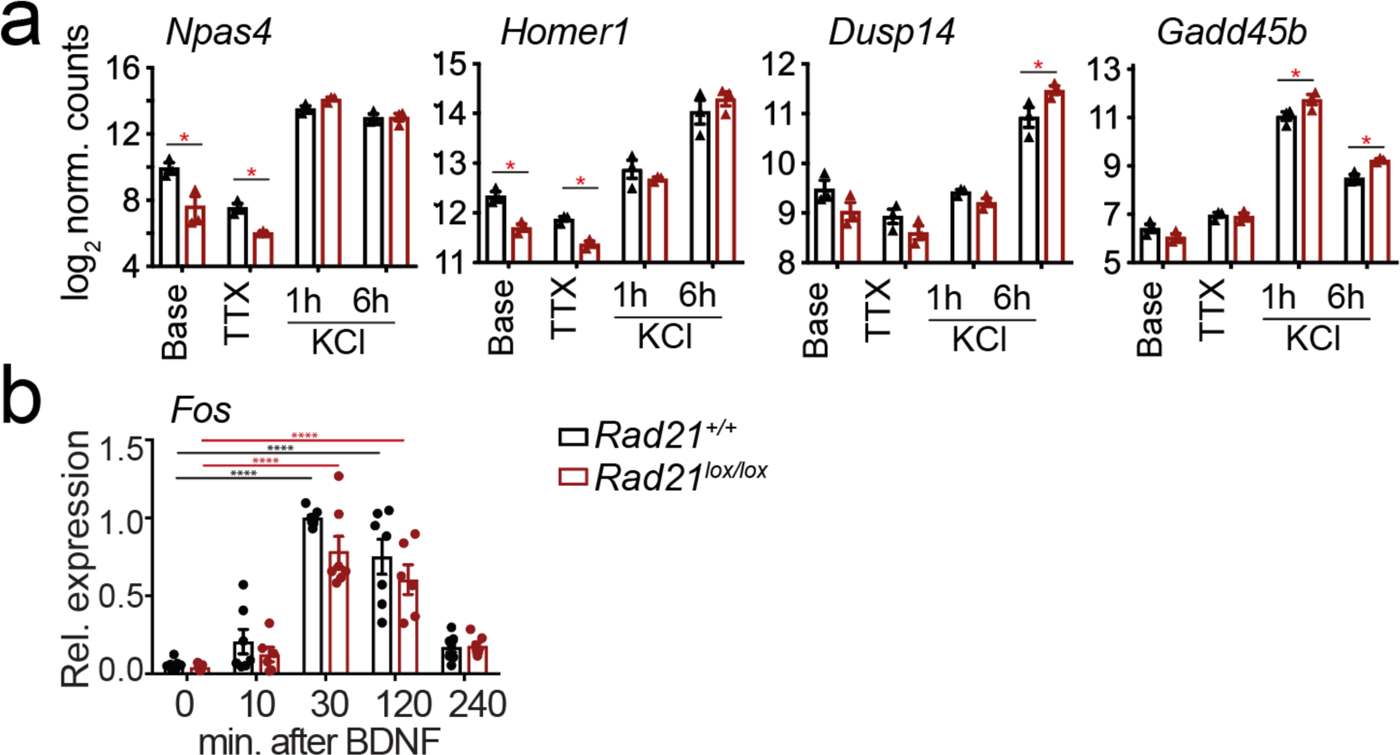
Inducible gene expression in cohesin-deficient neurons. a) Examples of ARG expression at baseline, after TTX/D-AP5 (TTX), and KCl-stimulation. Mean log2-transformed counts from 3 biological replicates (* adj. *P* < 0.05). b) Expression of *Fos* mRNA at baseline and at the indicated times after BDNF stimulation. Data points represent biological RT-PCR replicates. *P-*values refer to induction relative to 0 min. *** *P* <0.001.

**Supplementary Figure 9.**
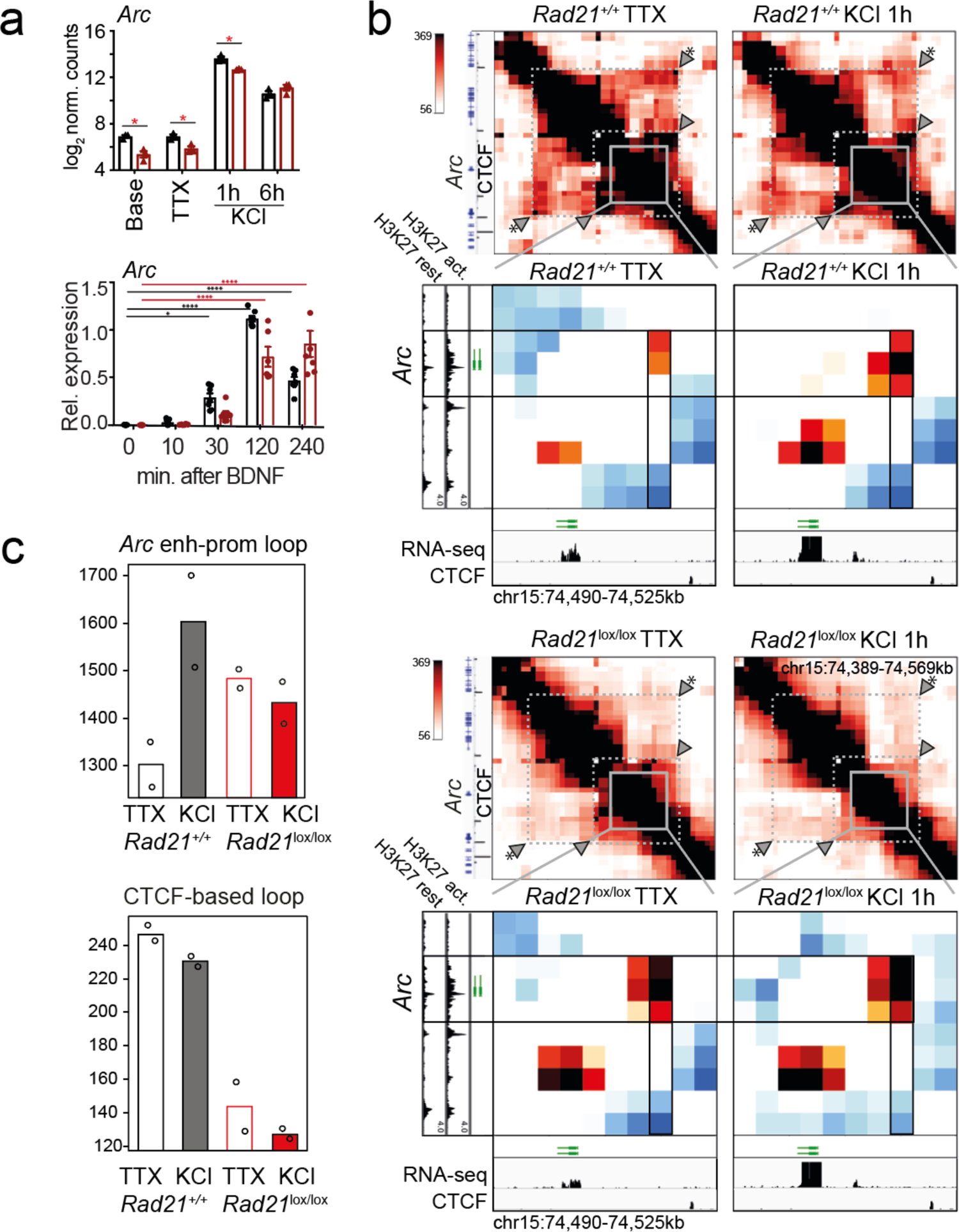
Contacts between the *Arc* promoter and an inducible enhancer in wild-type and cohesin-deficient neurons. a) Expression of *Arc* mRNA at baseline, after TTX/D-AP5 (TTX), and KCl-stimulation (top, mean log2-transformed counts from 3 biological RNA-seq replicates, * adj. *P* < 0.05) and at the indicated time after BDNF stimulation (bottom, data points represent biological RT-PCR replicates). *P-*values refer to induction relative to 0 min. * *P* < 0.05, *** *P* <0.001. b) Interaction score heatmaps of the ∼40 kb region immediately surrounding *Arc* obtained by 5C for resting (TTX) and 1h KCl-activated wild-type (top) and *Rad21*^lox/lox^ *Nex*^Cre^ neurons (bottom). Black frames highlight interaction between the *Arc* gene (y-axis) and a nearby downstream enhancer (x-axis). CTCF-ChIP-seq for cortical neurons is shown (Bonev et al., 2017). H3K27ac ChIP-seq in inactive (TTX-treated) and activated neurons is shown to annotate enhancer regions (Beagan et al., 2020). Two independent biological replicates are shown in Supplementary Fig. 10b. c) Quantification of 5C data. *Arc* enhancer-promoter loop (top). A CTCF-based loop that braces the *Arc* locus (arrowhead marked with * in panel b) is quantified for comparison (bottom).

**Supplementary Figure 10.**
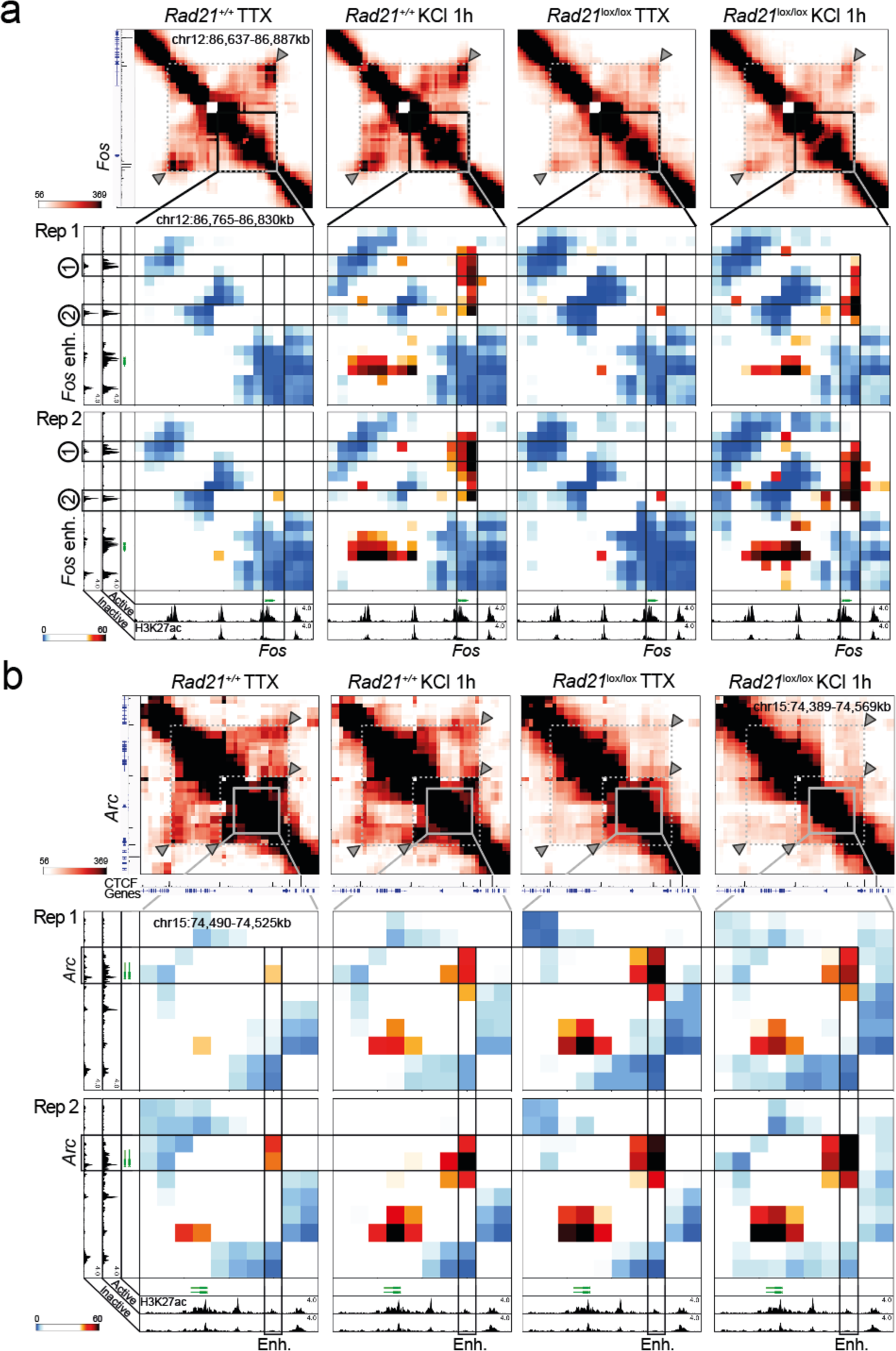
Replicate 5C cohesin depletion experiments. a) Top: Interaction frequency zoom-in heatmaps of 250 kb region surrounding the *Fos* gene. Dashed lines and arrow heads mark major CTCF binding sites at the boundaries of the domain that contains *Fos*. Note the weakening of these contacts in *Rad21*^lox/lox^ *Nex*^Cre^ neurons. Bottom: Interaction score heatmaps of the 65 kb region immediately surrounding the *Fos* gene. Black frames highlight interactions between the *Fos* gene and upstream enhancers 1 and 2. Two independent biological replicates are shown. H3K27ac ChIP-seq data (Beagan et al., 2020) from Bicuculline-treated (active) and TTX-treated (inactive) neurons annotate enhancer regions. b) Top: Interaction frequency zoom-in heatmaps of ∼200 kb region surrounding the *Arc* gene. Dashed lines and arrow heads mark major CTCF binding sites at the boundaries of domains that contain the Arc locus. Note the weakening of these contacts in *Rad21*^lox/lox^ *Nex*^Cre^ neurons. Bottom: Interaction score heatmaps of the ∼40 kb region immediately surrounding the *Arc* gene. Black frames highlight interaction between the *Arc* gene (y-axis) and a nearby downstream enhancer (x-axis). H3K27ac ChIP-seq data^25^ from Bicuculline-treated (active) and TTX-treated (inactive) neurons annotate enhancer regions. Two independent biological replicates are shown.

**Supplementary Figure 11.**
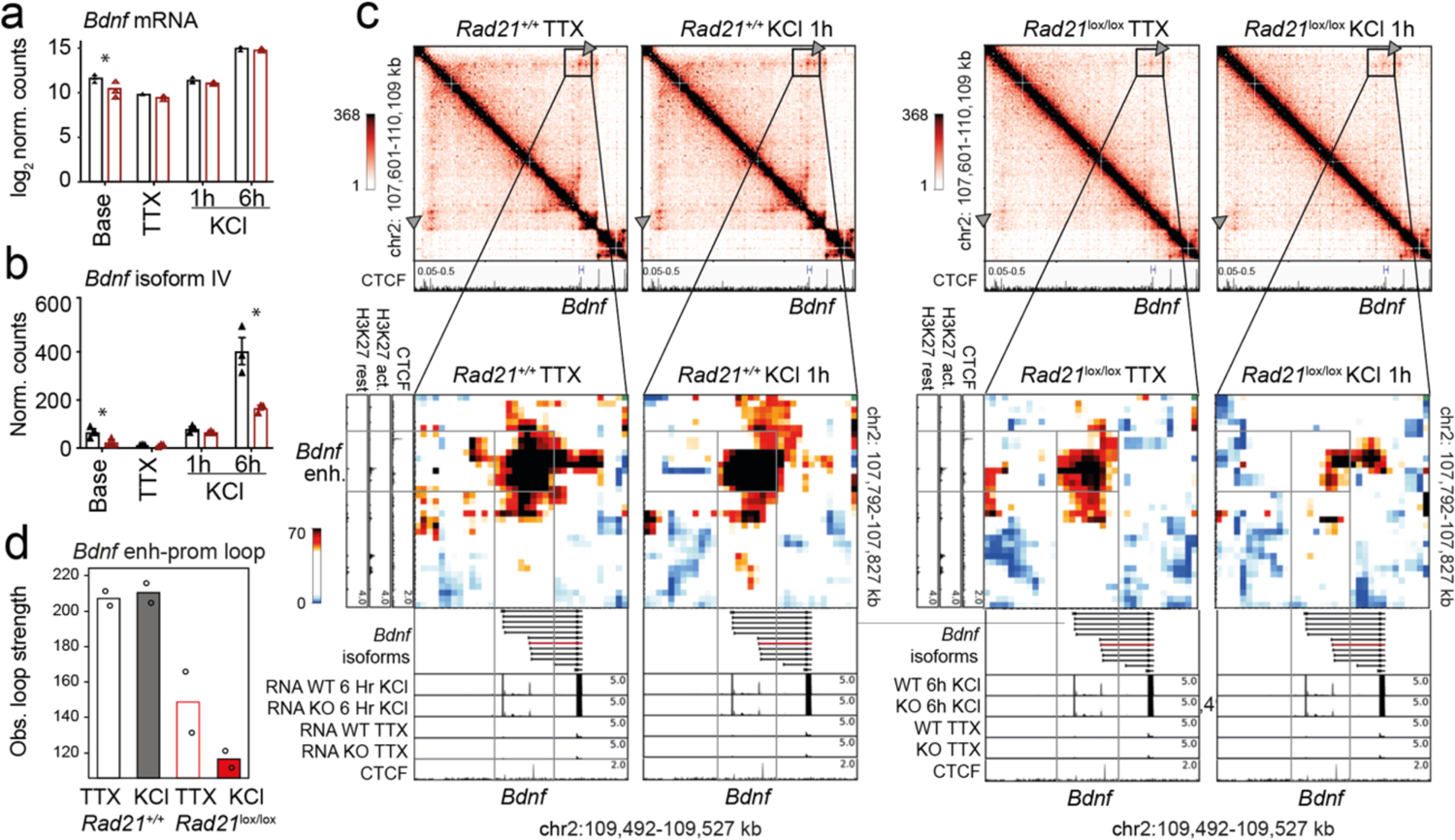
*Bdnf* enhancer-promoter IV contacts are weakend in cohesin-deficient neurons. a) Total *Bdnf* transcripts at baseline, after TTX/D-AP5 (TTX), and KCl-stimulation (left, mean log2-transformed counts from 3 biological replicates, * adj. *P* < 0.05). b) *Bdnf* promoter IV transcripts at baseline, after TTX/D-AP5 (TTX), and KCl-stimulation (left, mean log2-transformed counts from 3 biological replicates, * adj. *P* < 0.05). c) 5C interaction score heatmaps of the *Bdnf* region. CTCF ChIP-seq in cortical neurons (Bonev et al., 2017) and the position of *Bdnf* are displayed (top). Below: Zoom-in of constitutive *Bdnf* enhancer-promoter loop (gray frame). Shown on the side is H3K27ac ChIP-seq in resting and activated neurons, marking an activity-dependent enhancer, and CTCF ChIP-seq. RNA-seq in resting and activated wild-type and cohesin deficient neurons and CTCF ChIP-seq are shown underneath. Quantification of 5C contacts between *Bdnf* promoter IV and the activity-dependent enhancer.

## References

1. Aid T., Kazantseva A., Piirsoo M., Palm K., Timmusk T. Mouse and rat BDNF gene structure and expression revisited. J. Neurosci. Res., 85, 525–535 (2007)

2. Banerjee-Basu, S. & Packer, A. SFARI Gene: an evolving database for the autism research community. Dis. Model. Mech. 3, 133–135 (2010).

3. Beagan J, Phillips-Cremins JE. On the existence and functionality of topologically associating domains, Nature Genetics, 52, 8–16 (2020)

4. Beagan JA, Pastuzyn ED, Fernandez LR, Guo MH, Feng K, Titus KR, Chandrashekar H, Shepherd JD, Phillips-Cremins JE. Three-dimensional genome restructuring across timescales of activity-induced neuronal gene expression. Nat Neurosci. 23, 707–717 (2020)

5. Boija, A. et al. Transcription factors activate genes through the phase-separation capacity of their activation domains. Cell 175, 1842–1855 e16 (2018).

6. Bonev B, Mendelson Cohen N, Szabo Q, Fritsch L, Papadopoulos GL, Lubling Y, Xu X, Lv X, Hugnot JP, Tanay A, Cavalli G. Multiscale 3D Genome Rewiring during Mouse Neural Development. Cell 171: 557–572 (2017)

7. Cuartero, S. et al. Control of inducible gene expression links cohesin to hematopoietic progenitor self-renewal and differentiation. Nat. Immunol. 19, 932–941 (2018).

8. Crump NT, Ballabio E, Godfrey L, Thorne R, Repapi E, Kerry J, Tapia M, Hua P, Filippakopoulos P, Davies J. O. J., Milne T.A. BET inhibition disrupts transcription but retains enhancer-promoter contact BET inhibition disrupts transcription but retains enhancer-promoter contact. Nat Commun. 12, 223. doi: 10.1038/s41467-020-20400-z. (2021)

9. Deardorff, M. A., Noon, S. E. & Krantz, I. D. Cornelia de Lange Syndrome. In GeneReviews® (eds. Adam, M. P., et al.) (University of Washington, Seattle, 2018).

10. Dekker, J. & Mirny, L. The 3D Genome as Moderator of Chromosomal Communication. Cell 164, 1110–1121 (2016).

11. El Khattabi L, Zhao H, Kalchschmidt J, Young N, Jung S, Van Blerkom P, Kieffer-Kwon P, Kieffer-Kwon KR, Park S, Wang X, Krebs J, Tripathi S, Sakabe N, Sobreira DR, Huang SC, Rao SSP, Pruett N, Chauss D, Sadler E, Lopez A, Nóbrega MA, Aiden EL, Asturias FJ, Casellas R. A Pliable Mediator Acts as a Functional Rather Than an Architectural Bridge between Promoters and Enhancers. Cell 178:1145–1158.e20. (2019)

12. Fudenberg, G. et al. Formation of Chromosomal Domains by Loop Extrusion. Cell Rep. 15, 2038–2049 (2016).

13. Fujita, Y. et al. Decreased cohesin in the brain leads to defective synapse development and anxiety-related behavior. J Exp Med 214, 1431–1452 (2017)

14. Gallo, F.T., Katche, C., Morici, J.F., Medina, J.H., and Weisstaub, N.V. Immediate Early Genes, Memory and Psychiatric Disorders: Focus on c-Fos, Egr1 and Arc. Front Behav Neurosci 12, 79. (2018)

15. Goebbels, S. et al. Genetic targeting of principal neurons in neocortex and hippocampus of NEX-Cre mice. Genesis 44, 611–21 (2006).

16. Greer, P. L. & Greenberg, M. E. From synapse to nucleus: calcium-dependent gene transcription in the control of synapse development and function. Neuron 59, 846–60 (2008).

17. Gregor, A. et al. De novo mutations in the genome organizer CTCF cause intellectual disability. Am J Hum Genet 93, 124–31 (2013).

18. Greig, L. C., Woodworth, M. B., Galazo, M. J., Padmanabhan, H. & Macklis, J. D. Molecular logic of neocortical projection neuron specification, development and diversity. Nat. Rev. Neurosci. 14, 755–769 (2013).

19. Guo, Y. et al. CRISPR Inversion of CTCF Sites Alters Genome Topology and Enhancer/Promoter Function. Cell 162, 900–910 (2015).

20. Hirayama, T., Tarusawa, E., Yoshimura, Y., Galjart, N. & Yagi, T. CTCF is required for neural development and stochastic expression of clustered Pcdh genes in neurons. Cell Rep. 2, 345–57 (2012).

21. Hong EJ, McCord AE, Greenberg ME. A biological function for the neuronal activity-dependent component of Bdnf transcription in the development of cortical inhibition. Neuron 60: 610–24 (2008).

22. Hsieh T-H. S., Cattoglio C, Slobodyanyuk E, Hansen A.S., Xavier Darzacq X, Tjian R. Enhancer-promoter interactions and transcription are maintained upon acute loss of CTCF, cohesin, WAPL, and YY1. Biorxiv https://doi.org/10.1101/2021.07.14.452365 (2021)

23. Hsieh TS, Cattoglio C, Slobodyanyuk E, Hansen AS, Rando OJ, Tjian R, Darzacq X. Resolving the 3D Landscape of Transcription-Linked Mammalian Chromatin Folding. Mol Cell. 78: 539–553.e8 (2020)

24. Joo JY, Schaukowitch K, Farbiak L, Kilaru G, Kim TK. Stimulus-specific combinatorial functionality of neuronal c-fos enhancers. Nat Neurosci. 19, 75–83 (2016)

25. Kaech, S. & Banker, G. Culturing hippocampal neurons. Nat. Protoc. 1, 2406–15 (2006).

26. Kane L, Williamson I, Flyamer IM, Kumar Y, Hill RE, Lettice LA, Bickmore WA. Cohesin is required for long-range enhancer action. Biorxiv https://doi.org/10.1101/2021.06.24.449812 (2021)

27. Kawauchi, S. et al. Multiple organ system defects and transcriptional dysregulation in the Nipbl(+/-) mouse, a model of Cornelia de Lange Syndrome. PLoS Genet 5, e1000650 (2009).

28. Kim S, Yu NK, Shim KW, Kim JI, Kim H, Han DH, Choi JE, Lee SW, Choi DI, Kim MW, Lee DS, Lee K, Galjart N, Lee YS, Lee JH, Kaang BK. Remote Memory and Cortical Synaptic Plasticity Require Neuronal CCCTC-Binding Factor (CTCF). J Neurosci. 38: 5042–5052 (2018)

29. Kim, T. K. et al. Widespread transcription at neuronal activity-regulated enhancers. Nature 465, 182–7 (2010).

30. Krietenstein N, Abraham S, Venev SV, Abdennur N, Gibcus J, Hsieh TS, Parsi KM, Yang L, Maehr R, Mirny LA, Dekker J, Rando OJ. Ultrastructural Details of Mammalian Chromosome Architecture. Mol Cell. 78: 554–565.e7 c

31. Lupiáñez, D. G. et al. Disruptions of Topological Chromatin Domains Cause Pathogenic Rewiring of Gene-Enhancer Interactions. Cell 161, 1012–1025 (2015).

32. Malik, A. N. et al. Genome-wide identification and characterization of functional neuronal activity-dependent enhancers. Nat Neurosci. 17: 1330–9 (2014).

33. McCord, R., Kaplan, N. & Giorgetti, L. Chromosome Conformation Capture and Beyond: Toward an Integrative View of Chromosome Structure and Function. Molecular Cell 77, 688–708 (2020)

34. McGill BE, Barve RA, Maloney SE, Strickland A, Rensing N, Wang PL, Wong M, Head R, Wozniak DF, Milbrandt J. Abnormal Microglia and Enhanced Inflammation-Related Gene Transcription in Mice with Conditional Deletion of Ctcf in Camk2a-Cre-Expressing Neurons. J Neurosci. 38: 200–219 (2018)

35. Merkenschlager, M. & Nora, E. P. CTCF and Cohesin in Genome Folding and Transcriptional Gene Regulation. Annu. Rev. Genomics Hum. Genet. 17, 17–43 (2016).

36. Nasmyth, K. & Haering, C. H. Cohesin: its roles and mechanisms. Annu Rev Genet 43, 525– 58 (2009).

37. Nora, E. P. et al. Targeted Degradation of CTCF Decouples Local Insulation of Chromosome Domains from Genomic Compartmentalization. Cell 169, 930–944.e22 (2017).

38. Rajarajan, P., Gil, S. E., Brennand, K. J. & Akbarian, S. Spatial genome organization and cognition. Nat. Rev. Neurosci. 17, 681–691 (2016).

39. Rao, S. S. P. et al. A 3D map of the human genome at kilobase resolution reveals principles of chromatin looping. Cell 159, 1665–1680 (2014).

40. Rao, S. S. P. et al. Cohesin Loss Eliminates All Loop Domains. Cell 171, 305–320.e24 (2017).

41. Rinzema NJ, Sofiadis K, Tjalsma SJD, Verstegen MJAM Oz Y, aldes-Quezada C, Felder AK, Filipovska T, van der Elst S, de Andrade dos Ramos Z, Han R, Krijger PHL, de Laat W. Building regulatory landscapes: enhancer recruits cohesin to create contact domains, engage CTCF sites and activate distant genes. bioRxiv 10.1101/2021.10.05.463209 (2021)

42. Remeseiro, S., Cuadrado, A., Gomez-Lopez, G., Pisano, D. G. & Losada, A. A unique role of cohesin-SA1 in gene regulation and development. EMBO J. 31, 2090–102 (2012).

43. Rowley MJ, Corces VG. Organizational principles of 3D genome architecture. Nat Rev Genet. 19: 789–800 (2018)

44. Sabari, B. R. et al. Coactivator condensation at super-enhancers links phase separation and gene control. Science 361, eaar3958 (2018).

45. Sams, D. S. et al. Neuronal CTCF Is Necessary for Basal and Experience-Dependent Gene Regulation, Memory Formation, and Genomic Structure of BDNF and Arc. Cell Rep. 17, 2418–2430 (2016).

46. Sanz, E. et al. Cell-type-specific isolation of ribosome-associated mRNA from complex tissues. Proc Natl Acad Sci U A 106, 13939–44 (2009).

47. Schaukowitch, K. et al. Enhancer RNA facilitates NELF release from immediate early genes. Mol. Cell 56, 29–42 (2014).

48. Schwarzer, W. et al. Two independent modes of chromatin organization revealed by cohesin removal. Nature 551, 51–56 (2017).

49. Seitan, V. C. et al. A role for cohesin in T-cell-receptor rearrangement and thymocyte differentiation. Nature 476, 467–71 (2011).

50. Shin Y, Chang YC, Lee DSW, Berry J, Sanders DW, Ronceray P, Wingreen NS, Haataja M, Brangwynne CP. Liquid Nuclear Condensates Mechanically Sense and Restructure the Genome. Cell. 175: 1481–1491.e13 (2018)

51. Sholl, D. A. Dendritic organization in the neurons of the visual and motor cortices of the cat. J. Anat. 87, 387–406.1 (1953).

52. Spielmann, M., Lupiáñez, D. G. & Mundlos, S. Structural variation in the 3D genome. Nat. Rev. Genet. 19, 453–467 (2018).

53. Sun JH, Zhou L, Emerson DJ, Phyo SA, Titus KR, Gong W, Gilgenast TG, Beagan JA, Davidson BL, Flora Tassone, Phillips-Cremins JE. Disease-associated short tandem repeats co-localize to chromatin domain boundaries, Cell, 175, 224–238 (2018)

54. Thiecke MJ et al. Cohesin-Dependent and -Independent Mechanisms Mediate Chromosomal Contacts between Promoters and Enhancers. Cell Rep. 32:107929 (2020)

55. Tyssowski, et al. Different neuronal activity patterns induce different gene expression programs. Neuron. 98: 530–546.e11 (2018)

56. van den Berg, D. L. C., et al. Nipbl Interacts with Zfp609 and the Integrator Complex to Regulate Cortical Neuron Migration. Neuron 93, 348–361 (2017).

57. Weiss FD, Calderon L, Wang YF, Georgieva R, Guo Y, Cvetesic N, Kaur M, Dharmalingam G, Krantz ID, Lenhard B, Fisher AG, Merkenschlager M. Neuronal genes deregulated in Cornelia de Lange Syndrome respond to removal and re-expression of cohesin. Nat Commun. 12, 2919 (2021)

58. Won, H. et al. Chromosome conformation elucidates regulatory relationships in developing human brain. Nature 538, 523–527 (2016).

59. Yamada T, Yang Y, Valnegri P, Juric I, Abnousi A, Markwalter KH, Guthrie AN, Godec A, Oldenborg A, Hu M, Holy TE, Bonni A. Sensory experience remodels genome architecture in neural circuit to drive motor learning. Nature. 2019 May;569(7758):708-713.

60. Yap, E.L., and Greenberg, M.E. Activity-Regulated Transcription: Bridging the Gap between Neural Activity and Behavior. Neuron 100, 330–348.(2018)

## Supplementary references

61. Anders S, Pyl PT, Huber W. HTSeq--a Python framework to work with high-throughput sequencing data. Bioinformatics 31: 166–9 (2015).

62. Beaudoin, G. M. et al. Culturing pyramidal neurons from the early postnatal mouse hippocampus and cortex. Nat. Protoc. 7, 1741–54 (2012).

63. Durinck S, Spellman PT, Birney E, Huber W. Mapping identifiers for the integration of genomic datasets with the R/Bioconductor package biomaRt. Nat Protoc. 4: 1184–91 (2009).

64. Fernandez LR, Thomas G., Gilgenast TG, Phillips-Cremins JE. 3DeFDR: statistical methods for identifying cell type-specific looping interactions in 5C and Hi-C data, Genome Biology, 21;219, (2020)

65. Gao F, Cam HP, Tsai L-H. Neutools: A collection of bioinformatics web tools for neurogenomic analysis. bioRxiv 429639; doi: https://doi.org/10.1101/429639a (2018)

66. Gilgenast, T. G. & Phillips-Cremins, J. E. Systematic evaluation of statistical methods for identifying looping interactions in 5C data. Cell Syst. 8, 197–211 e113 (2019)

67. Imakaev, M. et al. Iterative correction of Hi-C data reveals hallmarks of chromosome organization. Nat. Methods 9, 999–1003 (2012).

68. Kim D, Pertea G, Trapnell C, Pimentel H, Kelley R, Salzberg SL. TopHat2: accurate alignment of transcriptomes in the presence of insertions, deletions and gene fusions. Genome Biol. 14(4): R36 (2013).

69. Kim JH, Titus KR, Gong W, Beagan JA, Cao Z, Phillips-Cremins JE, 5C-ID: Increased resolution Chromosome-Conformation-Capture-Carbon-Copy with in situ 3C and double alternating primer design, Methods, 142: 39–46, 2018

70. Lajoie, B. R., van Berkum, N. L., Sanyal, A. & Dekker, J. My5C: web tools for chromosome conformation capture studies. Nat. Methods 6, 690–691 (2009).

71. Langmead B, Trapnell C, Pop M, Salzberg SL. Ultrafast and memory-efficient alignment of short DNA sequences to the human genome. Genome Biol. 10(3): R25 (2009).

72. Lawrence M, Huber W, Pagès H, Aboyoun P, Carlson M, Gentleman R, Morgan MT, Carey VJ. Software for computing and annotating genomic ranges. PLoS Comput Biol. 9: e1003118 (2013)

73. Love, M. I., Huber, W. & Anders, S. Moderated estimation of fold change and dispersion for RNA-seq data with DESeq2. Genome Biol. 15, 550 (2014).

74. Nelson JD, Denisenko O, Bomsztyk K. Protocol for the fast chromatin immunoprecipitation (ChIP) method. Nat Protoc. 1, 179–85 (2006).

75. Paxinos G, Halliday G, Watson C, Kassem M. Atlas of the Developing Mouse Brain. 2nd Edition. Academic Press, 2020

76. Su, Y. et al. Neuronal activity modifies the chromatin accessibility landscape in the adult brain. Nat. Neurosci. 20, 476–483 (2017).

77. Subramanian, A. et al. Gene set enrichment analysis: a knowledge-based approach for interpreting genome-wide expression profiles. Proc Natl Acad Sci U A 102, 15545–50 (2005)

78. Tachibana-Konwalski, K. et al. Rec8-containing cohesin maintains bivalents without turnover during the growing phase of mouse oocytes. Genes Dev 24, 2505–16 (2010).

79. Young, M. D., Wakefield, M. J., Smyth, G. K. & Oshlack, A. Gene ontology analysis for RNA-seq: accounting for selection bias. Genome Biol. 11, R14 (2010).

80. Zhang Y. et al. Model-based analysis of ChIP-Seq (MACS). Genome Biol. 9: R137 (2008).

